# The Cdc42 effector and non-receptor tyrosine kinase, ACK defines a distinct STAT5 transcription signature in a chronic myeloid leukaemia cell model

**DOI:** 10.1101/2022.06.10.494364

**Authors:** Jessica Corry, Daniel Jachimowicz, Benjamin Keith, Jose Julio Vicente-Garcia, Helen R. Mott, Kate Wickson, Darerca Owen

## Abstract

Activated Cdc42-associated kinase (ACK) is a Rho family effector that is widely implicated in cancer. Here, we describe new roles for ACK in transcriptional regulation mediated by its relationship with the signal transducer and activators of transcription (STAT) family. We show that ACK can interact with STAT3, STAT5A and STAT5B, and augments phosphorylation at the conserved activation tyrosine on these STAT members. ACK stimulates oncogenic STAT nuclear relocation and transcriptional activation. We also identify endogenous relationships between ACK and STAT family members in haematopoietic disease cell lines. In the K562 chronic myeloid leukaemia cell line, we confirm that ACK contributes to the pool of active, nuclear STAT5. By interrogating ACK knock out cells we describe an ACK-driven STAT5 transcriptional signature in K562s. We propose ACK as a contributor to hyperactivated STAT5 signalling in this CML cell line and reveal a new route for therapeutic intervention.

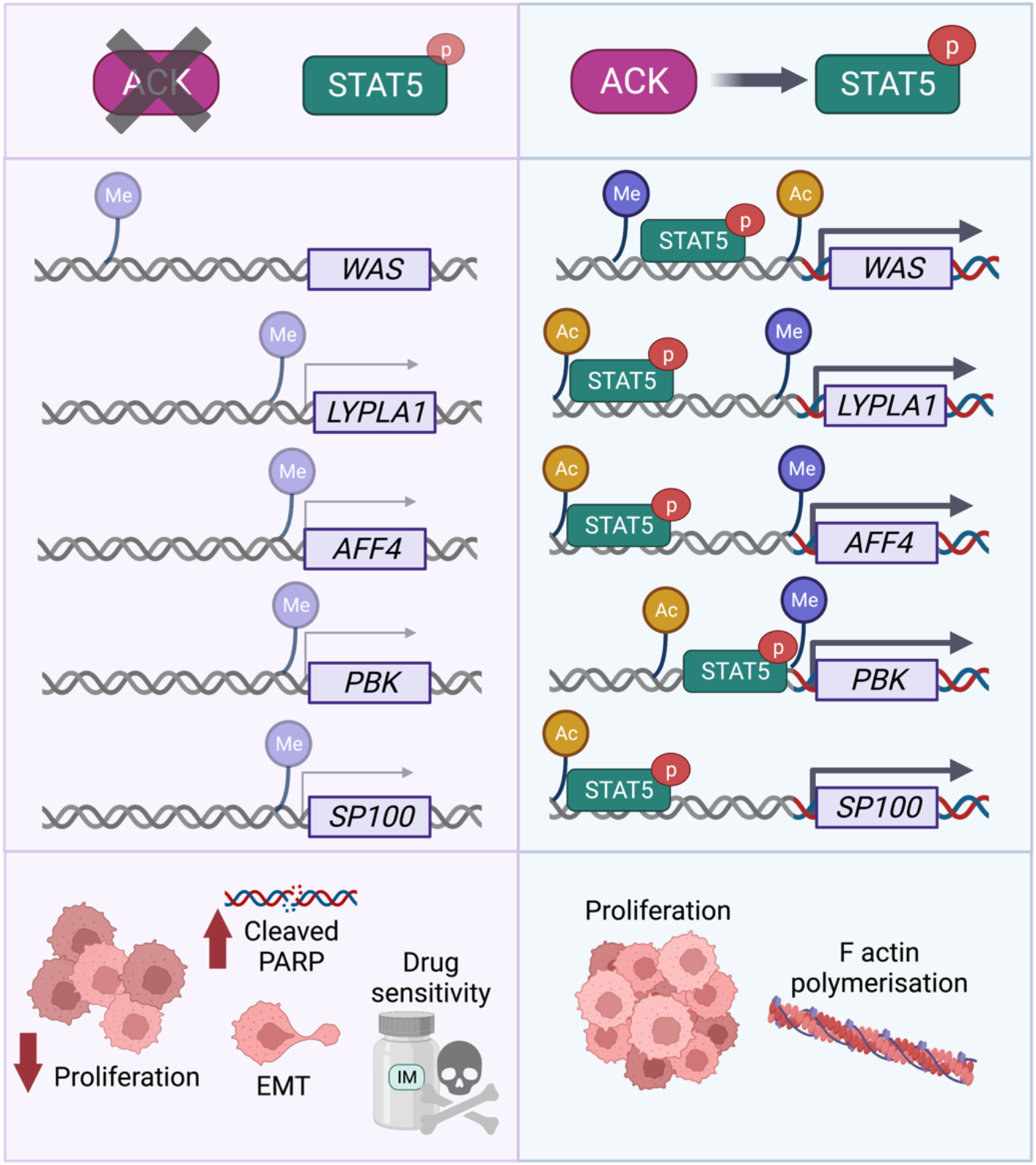

## Introduction

Tyrosine kinases regulate intracellular signalling pathways that are critical for both health and disease. By acting as critical mediators of protein activation and deactivation, they serve as core regulators of signalling pathways and are attractive therapeutic targets. In the two decades since the FDA approval of imatinib, the first kinase targeting drug, considerable progress in the kinase inhibitor field has taken place. However, for many kinases, a refined understanding of the molecular pathways they govern is unavailable and will be crucial for successful drug targeting.

One important non-receptor tyrosine kinase (NRTK) that is widely implicated in cancer, yet not currently pharmacologically inhibited in the clinic, is activated Cdc42 associated kinase (ACK). ACK is encoded by *TNK2* on chromosome 3q29, a region amplified in a plethora of cancer types (Van de Horst et al., 2005). Such amplification results in elevated ACK expression which is often correlated with poor prognosis (Fox, Crafter and Owen, 2019). The roles of ACK in both solid state and haematopoietic disease have been recently reviewed (Wang et al., 2021), describing the involvement of ACK in leukaemia, melanoma, prostate, gastric, lung, liver and breast cancer. In addition to many ACK somatic mutations being identified in cancer samples, ACK has more importantly been highlighted as a kinase likely to carry a cancer driver mutation (Greenman et al., 2007; Prieto-Echangüe et al., 2010).

ACK is a ubiquitously expressed kinase, which was initially discovered as an effector of the small G protein Cdc42, interacting with Cdc42·GTP via its Cdc42/Rac binding (CRIB) region (Manser et al., 1993). ACK contains multiple domains and functional regions including: a sterile alpha motif (SAM) domain, a nuclear export signal (NES), a kinase domain, a Src homology 3 (SH3) domain, a proline rich region, a clathrin binding domain and a ubiquitin association domain (UBA). With this array of domains, ACK can engage a wide range of partners including the tumour suppressor WWOX, the androgen receptor (AR), the epidermal growth factor receptor (EGFR), Wiskott-Aldrich syndrome protein (WASP), phosphoinositide 3 kinase (PI3K), protein kinase B (AKT), growth factor receptor bound protein 2 (Grb2), cortactin, clathrin and ubiquitin (Fox, Crafter and Owen, 2019).

ACK has also been shown to influence the transcriptional landscape in hormone responsive cancer models by functioning as both an epigenetic modifier and transcriptional co-activator. In prostate cancer cell lines, ACK was shown to bind, phosphorylate and activate the AR, with the ACK-AR complex then binding upstream of target genes including prostate specific antigen (PSA) (Mahajan et al., 2007). In this setting therefore, ACK acts as a transcriptional co-activator, influencing AR transcriptional output. Additionally, ACK is known to influence epigenetic changes that control the transcription of the AR itself (Mahajan et al., 2017). In castration resistant prostate cancer (CRPC) cell lines, ACK was shown to interact with and phosphorylate Tyr88 of histone 4, which then acted as a transcription activating mark. ChIP-seq in a CRPC cell line identified these marks upstream of the AR transcriptional start site (TSS), marking the site for transcription. Inhibition or genetic deletion of ACK ablated these epigenetic marks, along with AR expression. Small molecule inhibition of ACK has also been shown to significantly reduce AR mRNA expression in tissues and reduce *in vivo* CRPC tumour growth (Mahajan et al., 2017). ACK also influences epigenetic aspects of oestrogen receptor (ER)-driven transcription in breast cancer models (Mahajan et al., 2014). In MCF7 cells, ACK has been shown to bind, phosphorylate and therefore activate the H3K9 histone demethylase, KDM3A. HOXA1, an ER target gene, has increased H3K9me levels at its TSS in ACK inhibited MCF7 cells, due to the altered activation of KDM3A, which results in downregulated HOXA1 expression (Mahajan et al., 2014). Thus, ACK can bind to key transcription factors, modify histones and alter the activation of pivotal epigenetic regulators to influence transcription in both breast and prostate cancer models.

Additional transcriptional proteins which interact with ACK include STAT1 and STAT3, members of the signal transducer and activators of transcription (STAT) family (Mahendrarajah et al., 2017). The STAT family are master regulators of gene transcription and include three key oncogenic members: STAT3, STAT5A and STAT5B (Buettner, Mora and Jove, 2002). Critical to STAT activation is phosphorylation of a conserved tyrosine residue, which facilitates dimerization by supporting an interaction with an SH2 domain of a second STAT monomer. Despite being so widely implicated in cancer, no direct STAT inhibitor has yet made it to the clinic. Instead, research efforts have focussed on the hyperactivated STAT activators as tractable targets. A previous yeast-2-hybrid screen in our lab (Clayton et al., 2022) identified STAT3 as a potential ACK interacting partner, a relationship that has been reported in the literature (Mahendrarajah et al., 2017). In this study we confirm ACK as a STAT3 interacting partner and present, for the first time, relationships between ACK, STAT5A and STAT5B. We explore the role of this interaction within a chronic myeloid leukaemia (CML) cell line and identify changes to the STAT5 transcriptional network upon loss of ACK in K562 cells, defining an ACK-driven STAT5 transcriptional signature in the CML cell model.

## Results

### ACK interacts with the oncogenic transcription factors STAT3 and STAT5 in haematopoietic cell models

We first sought to validate that STAT3 is an ACK interacting partner in human cells by co-immunoprecipitation. Wild type (wt) ACK or K158R, a kinase dead mutant ACK (dACK), were transiently expressed alone or with FLAG-STAT3 in HEK293T cells and subjected to FLAG immunoprecipitation. Immunoblots confirmed a kinase dependent interaction between wtACK and STAT3 (Figure 1A). A strong increase in the phosphorylation of Tyr705-STAT3 was also observed upon ACK co-expression. Tyr705 is strictly conserved between STAT family members and its phosphorylation is critical for their activation (Shuai et al., 1992). Since a constitutively active ACK variant has previously been shown to interact with STAT1 (Mahendrarajah et al., 2017), we investigated if other STAT family members were ACK partners. Co-immunoprecipitation also confirmed an interaction between wtACK and both STAT5A and STAT5B, along with clear enhancement of tyrosine phosphorylation at the conserved Tyr694/699 site (Figure 1A). We next confirmed that the observed phosphorylation stimulated STAT transcriptional activity using a luciferase reporter assay. STAT3, STAT5A and STAT5B were transiently expressed in HEK293T cells alongside a reporter and *Renilla* transfection control, individually and in combination with wtACK. Protein expression was confirmed by immunoblot (Figure 1B). Co-expression with wtACK induced a significant increase in luciferase levels with all three STATs, indicating increased transcriptional activity (Figure 1B). Transcriptional activation of the STATs was also inhibited by AIM-100, an ACK inhibitor. Thus, ACK interacts with all three oncogenic STATs, phosphorylates a critical tyrosine residue and stimulates their transcriptional activity.

**Figure 1:**
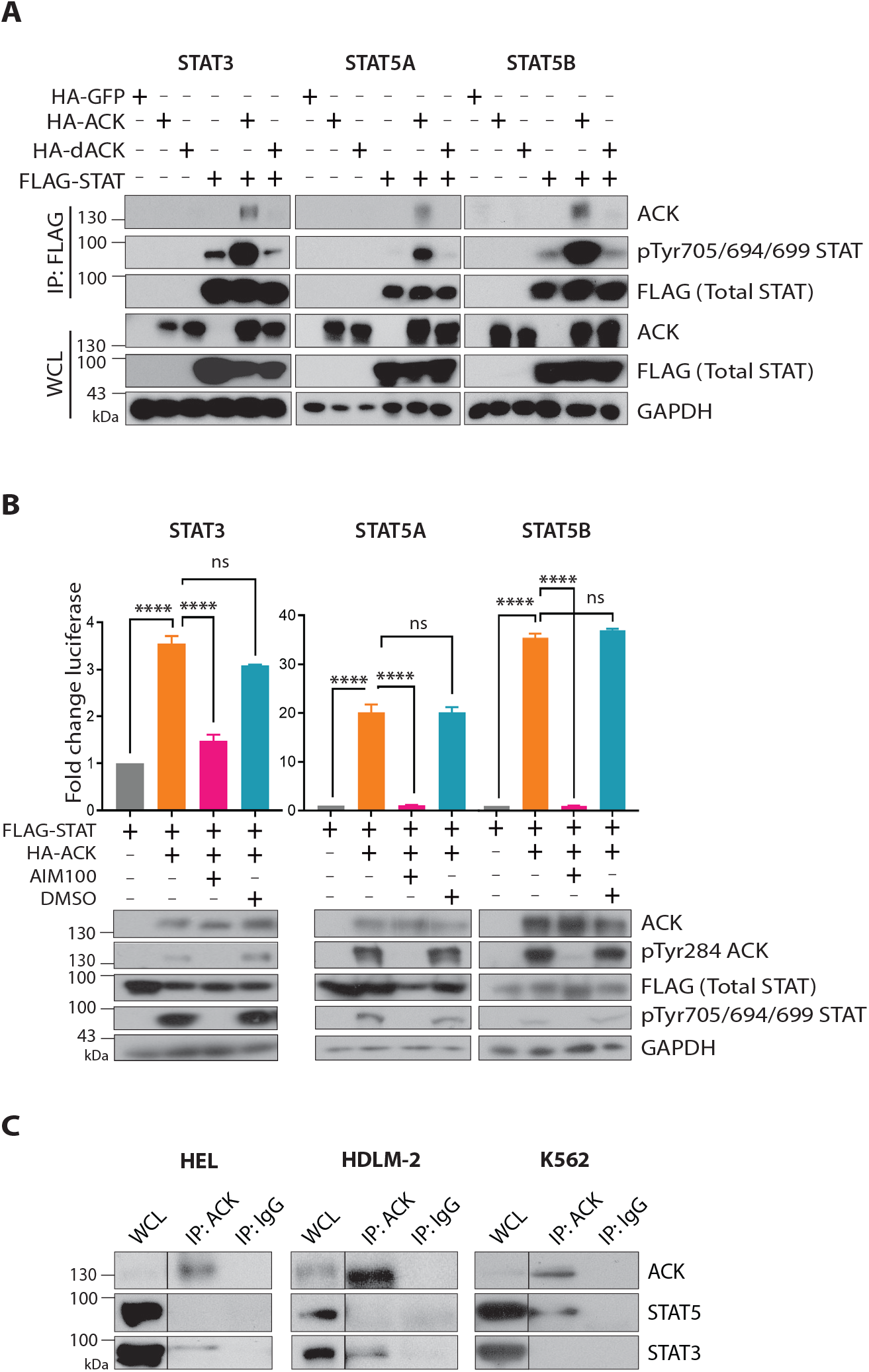
ACK interacts with STAT3, STAT5A and STAT5B and stimulates transcriptional activity. **(A)** Co-immunoprecipitation of ACK and STAT family members. HA-GFP, HA-ACK or HA-dACK (kinase dead mutant) were expressed alone or in combination with FLAG-STAT in HEK293T cells. Cells were lysed, subjected to anti-FLAG immunoprecipitation and analysed by western blot. GAPDH is a loading control. WCL=whole cell lysate. N=3. **(B)** Luciferase reporter assays showing STAT transcriptional activity. FLAG-STAT, STAT luciferase reporter and *Renilla* expression control were transiently expressed in HEK293T cells with empty vector or HA-ACK, and if stated 10 µM (STAT3) or 1 µM (STAT5A/B) of the ACK inhibitor AIM-100/DMSO 16 hours pre lysis. Luciferase signals were normalized to *Renilla* and are presented as fold change in signal with respect to cells expressing STAT alone. Significance was determined with a one-way ANOVA followed by Tukey’s multiple comparison test where ns=not significant, **** = p<0.0001. N=3, error bars show SEM. Protein expression in lysates was confirmed by immunoblot. GAPDH is a loading control. **(C)** Immunoprecipitation of endogenous ACK in human cancer cell lines. K562, HEL and HDLM-2 cells were lysed and subject to overnight immunoprecipitation with anti-ACK or an IgG isotype control. Samples were analysed by immunoblot to confirm endogenous interactions with STAT family members. N=3.

We next canvassed a panel of cancer cell lines (Table S1) to identify endogenous ACK/STAT interactions. Although we were unable to detect an interaction in the solid-state cell models tested, complexes were observed in several haematopoietic disease cell lines. Endogenous ACK was confirmed to interact with STAT3 in the human erythroleukaemia line, HEL and the Hodgkin’s lymphoma line, HDLM-2 (Figure 1C). Additionally, ACK was shown to interact with STAT5 in the CML cell line, K562 (Figure 1C). To our knowledge, these are the first reports of endogenous ACK-STAT interactions and indicate a possible specificity of the ACK-STAT relationship in haematopoietic disease models.

### ACK increases levels of active, nuclear STAT

Classical STAT activation involves cytoplasmic phosphorylation, dimerization and nuclear translocation (Corry, Mott and Owen, 2020). To investigate if the cellular localization of oncogenic STATs is altered upon co-expression with ACK, HEK293T cells were separated into cytoplasmic and nuclear-enriched fractions. Upon co-expression with wtACK, a significant increase in the percentage of total STAT3, STAT5A and STAT5B was observed in the nuclear-enriched fraction (Figure 2A, lower panels and B). Phosphorylated STATs were observed in both fractions (Figure 2A), suggesting that STAT activation occurs in accordance with the canonical model of STAT signalling (Corry, Mott and Owen, 2020). STAT activation was also monitored using confocal microscopy. Co-expression of ACK with STAT3, STAT5A and STAT5B in HEK293T/17 cells induced strong STAT phosphorylation (Figure 2C-E). Columbus image analysis software was used to confirm that there was a significant increase in the percentage of total cells exhibiting the pTyr-STAT signal upon ACK co-expression (Figure 2C-E). Thus, upon ACK co-expression in human cells, a clear increase in the nuclear presence of active, phosphorylated STAT is observed.

**Figure 2:**
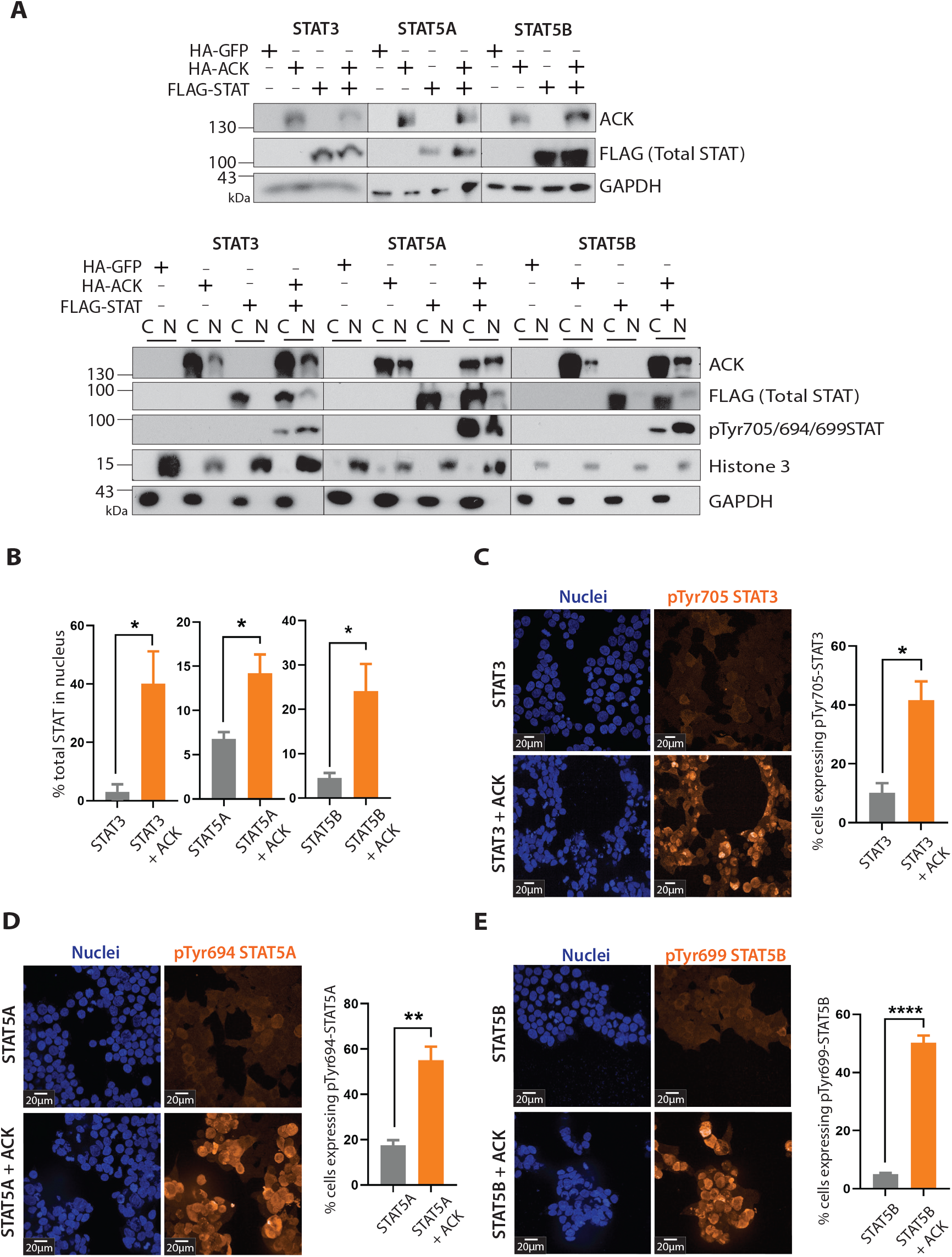
Nuclear localization of phosphorylated STAT. **(A)** Biochemical fractionation to monitor cellular localization changes of STATs. HEK293T cells were transiently transfected with HA-GFP, HA-ACK alone or in combination with FLAG-STAT and subject to biochemical fractionation into nuclear (N) and cytoplasmic (C) enriched fractions (lower panel). Total protein expression before and after fractionation was confirmed by immunoblot (top panel). GAPDH and Histone 3 were used as cytoplasmic and nuclear markers respectively. N=3. **(B)** Densitometry of fractionation immunoblots. N=3, error bars show SEM. Significance determined by unpaired student t-test where * p <0.05. **(C-E)** Confocal microscopy of HEK293T/17 cells transiently expressing FLAG-STAT alone or in combination with HA-ACK. Images were obtained using x60 water magnification lens. Scale bar of 20 µM shown. Nuclei were stained with Hoescht (blue). The orange signal represents pTyr705-STAT3, pTyr694-STAT5A or pTyr699-STAT5B. Columbus image analysis was used to count the number of cells expressing active STAT upon ACK co-expression. N=3, error bars show SEM. Significance determined by unpaired student t-test where * p<0.05, ** p<0.01 and *** p<0.001.

### The ACK-STAT5 axis in K562 chronic myeloid leukaemia cells

We have shown an interaction between ACK and STAT5 in the K562 CML cell line (Figure 1C). It has also been reported that K562s treated with histone deacetylase complex inhibitors (HDACi) show caspase-induced ACK degradation and loss of pTyr705-STAT3 (Mahendrarajah, Paulus and Krämer, 2016). To further investigate the ACK-STAT relationship in the K562s we generated ACK KO clonal cell lines using CRISPR-Cas9 RNP based targeting. Immunoblot analysis of whole cell lysates from three KO cell lines confirmed ablation of ACK expression as well as significantly reduced pTyr-STAT5, with no change to total STAT5 levels (Figure 3A and B). We saw no effect on STAT3 phosphorylation in ACK KO cells (Figure S1) in accordance with the lack of an ACK-STAT3 interaction observed in this cell line (Figure 1C). The reduction in endogenous pTyr-STAT5 in ACK KO cells was also assessed by confocal microscopy, with a significant reduction in cellular pTyr-STAT5 confirmed by image analysis (Figure 3C). Since pTyr-STAT5 is a known proliferative driver in K562 cells (Weber et al., 2015) we tested whether loss of ACK, and subsequent pTyr-STAT5, affected cell growth and found that ACK KO resulted in a significant reduction in cell proliferation (Figure 3D). To further investigate the localization of the remaining pTyr-STAT in ACK KO cells, we subjected cells to biochemical fractionation. This revealed a significant loss of phosphorylated STAT5 in the nuclear-enriched fraction in two out of three ACK KO clones, although the overall trend was evident in all (Figure 3E and F). Thus, ACK regulates, at least in part, the nuclear pool of phosphorylated and transcriptionally active STAT5 in K562 cells, which leads to a disruption to STAT5 signalling and reduced proliferation in ACK KO K562 cells.

**Figure 3:**
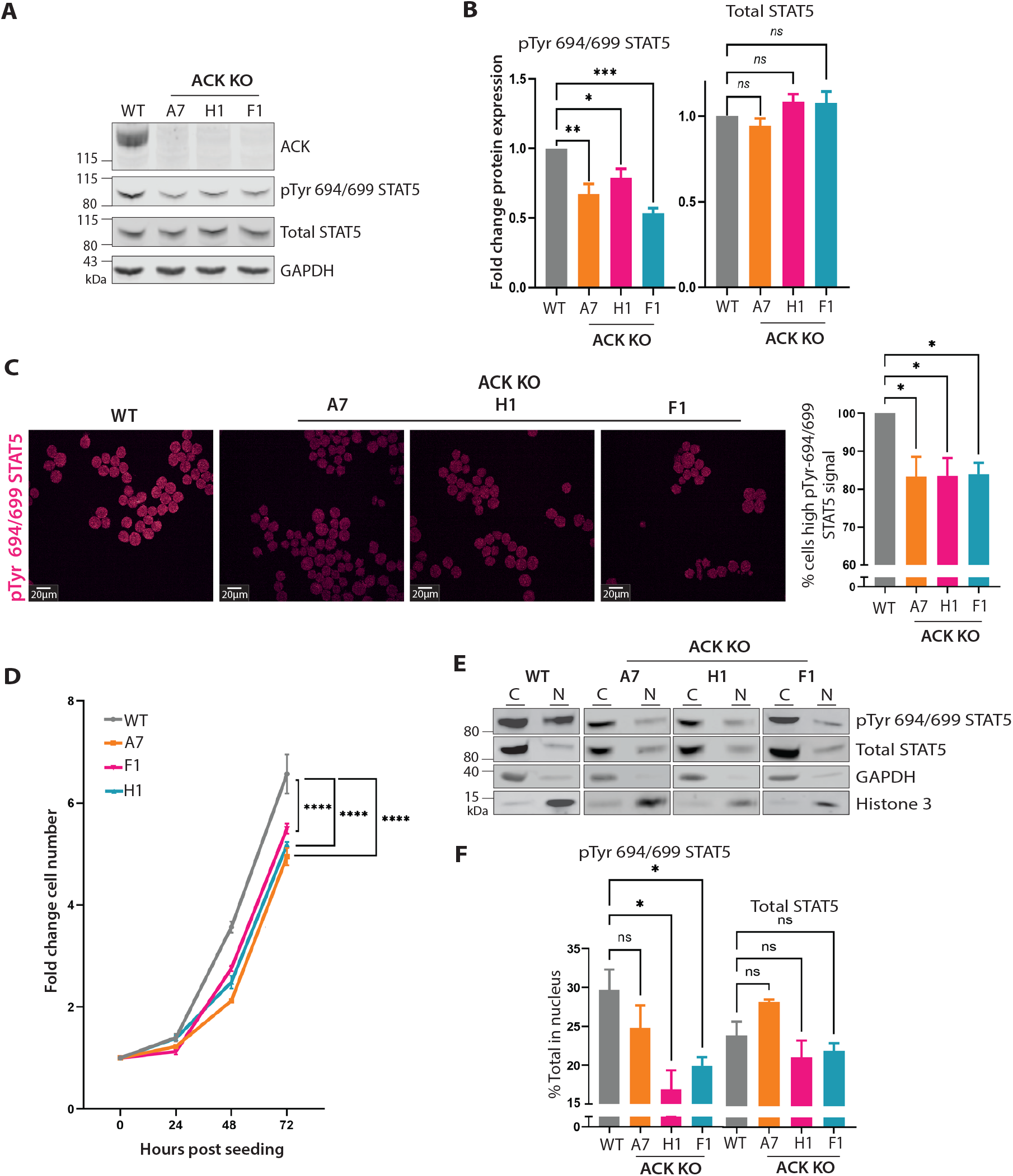
ACK-STAT5 relationship in a chronic myeloid leukaemia cell line. **(A)** Generation of ACK KO K562 clonal cell lines by CRISPR-Cas9 RNP based targeting. Protein expression was assessed in lysates by immunoblot. GAPDH is a loading control. N=3. **(B)** Quantification of total STAT5 and pTyr694STAT5A/pTyr699STAT5B levels in ACK KO cells by densitometry. N=3, error bars show SEM. Statistical significance determined by one-way ANOVA followed by Dunnett’s multiple comparisons test, with each condition compared to WT, ns=not significant, * p<0.05, ** p <0.01 and *** p <0.001. **(C)** Confocal microscopy of endogenous pTyr694STAT5A/pTyr699STAT5B in K562 cells, scale bars of 20 µM shown. Images obtained using x60 water magnification lens. RHS: Columbus image analysis was used to count the number of cells expressing pTyr 694STAT5A/pTyr699STAT5B over a threshold. N=3, error bars show SEM. Statistical significance determined by one-way ANOVA followed by Dunnett’s multiple correction test, with each condition compared to WT, * p<0.05. **(D)** 0.5 million K562 cells were seeded in 12 well plates and incubated over 72 hours. Every 24 hours, cell number was determined using a Beckman automated cell counter. Data from 3 combined biological repeats are shown. Statistical significance determined by a two-way ANOVA followed by Dunnett’s multiple correction test. *** p=0.001, **** p<0.001. **(E)** Cellular distribution of total STAT5 and pTyr694STAT5A/pTyr699STAT5B in WT and ACK KO K562 was assessed by biochemical fractionation into cytoplasmic (C) and nuclear (N) enriched fractions, and determined by immunoblot. GAPDH and Histone 3 were used as cytoplasmic and nuclear markers respectively. N=3. **(F)** Quantification of pTyr694STAT5A/pTyr699STATB and total STAT5 levels in the nucleus by densitometry. N=3, error bars show SEM. Statistical significance determined by one-way ANOVA followed by Dunnett’s multiple comparisons test, with each condition compared to WT, * p<0.05.

### ACK-directed STAT5 transcriptional control

To understand how the STAT5 transcriptional network is altered in ACK KO K562s, we first sought to identify changes in STAT5 DNA binding. We performed <u>c</u>leavage <u>u</u>nder <u>t</u>argets and <u>r</u>elease <u>u</u>sing <u>n</u>uclease (CUT&RUN) (Skene and Henikoff, 2017) against STAT5 using the F1 ACK KO clone. We first determined the number of consensus peaks in the CUT&RUN data set between replicates, where peaks indicate regions of STAT5 binding. In WT cells a total of 747 consensus STAT5 peaks were identified, compared to 418 in the ACK KO, confirming a global loss of STAT5 DNA binding (Figure 4A, left). In addition, we also investigated histone methylation (H3K4me3) and histone acetylation (H3K27ac) as markers of active promoter and enhancer regions (Figure 4A, centre and right). Global loss of both histone marks was observed in ACK KO cells, confirming that the histone landscape is altered in ACK KO cells, with the greatest reduction seen in active enhancer marks.

**Figure 4:**
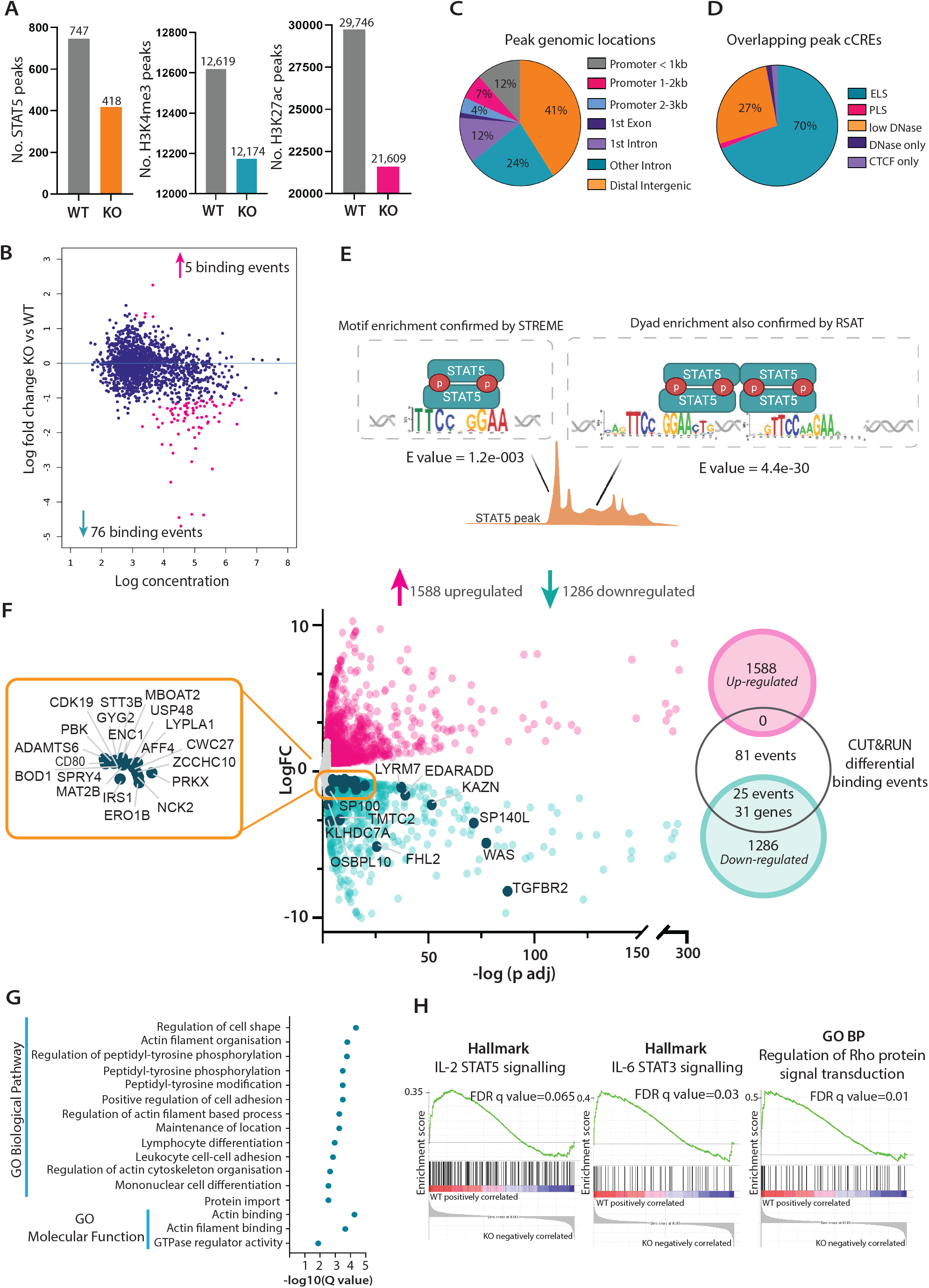
ACK regulated STAT5 DNA binding in K562 cells. **(A)** STAT5 DNA binding was assessed by CUT&RUN. Consensus peak numbers called for STAT5, H3K4me3 and H3K27ac in WT and ACK KO F1 clone are represented in bar charts. **(B)** 81 differential STAT5 binding events in ACK KO cells identified by DiffBind and represented as an MA plot. MA plots are presented with concentration on the X-axis, reflective of the mean normalised number of reads across replicates for a binding site, compared to the fold change of STAT binding at the locus (Y-axis). Pink dots represent hits with an FDR (false discovery rate) <0.05. **(C)** Genomic location of differential binding events defined by ChIPseeker. **(D)** Overlap of differential binding events and ENCODE candidate *cis* regulatory elements (cCRE). cCREs classed as promoter like signature (PLS), enhancer like signature (ELS), CCCTC binding factor (CTCF) and low or standard sensitivity to DNase (reflective of open chromatin) signatures. **(E)** Graphic to demonstrate different *in silico* analysis tools to identify STAT motif enrichment that would allow for dimer binding (by STREME) and potential tetramer binding (by RSAT). **(F)** Differential gene expression between WT and F1 ACK KO K562. Pink dots represent the 1588 upregulated genes and turquoise dots the 1286 downregulated genes. Genes that were also identified by GREAT as under regulation of the lost STAT5 binding peaks are labelled in the orange box. The x-axis represents -log of adjusted p values calculated by DEseq2 vs the log fold change (FC) on the Y-axis. Venn diagram to represent the overlap between the GREAT assigned genes from the differential binding events and differential expression. None of the 5 gained binding events could be linked to an upregulated gene. 25 binding events lost in KO cells could be linked to 31 upregulated genes. **(G)** OncoenrichR plot of GO biological pathway and GO molecular function pathways (blue bars) that are enriched in WT cells. Q value (graph, RHS) represents the p value for each gene set, adjusted for the false discovery rate. **(H)** GSEA outputs. A peak to the LHS demonstrates enrichment of named gene sets (BP=Biological Process) in WT cells, and thus pathways disrupted in ACK KO cells.

To identify locations of differential STAT5 binding between WT and KO cells we next used DiffBind. This identified 81 statistically significant, differentially bound sites, with the majority (76) being losses in STAT5 binding in KO cells (Figure 4B). Strong correlation was observed between biological replicates of each condition (WT vs KO) (Figure S2A). ChIPseeker was then used to annotate the genomic location of each differentially bound peak. This analysis showed that 41% of differentially bound peaks were intergenic, 35% intronic and 22% within 3kb of a promoter (Figure 4C). The large percentages of peaks observed in promoter-distal locations correlated with the large loss in global histone acetylation seen in the KO cells. Additionally, we analyzed whether the lost peak locations overlapped with candidate cis-regulatory elements (cCREs) identified in K562s (The ENCODE project consortium et al., 2020). cCREs were identified to overlap with the 76 lost STAT5 binding regions and, overwhelmingly, 70% were of an enhancer-like signature (ELS) (Figure 4D), thus confirming the importance of ACK in STAT5 enhancer binding in K562 cells.

STAT family members bind to DNA at a conserved binding motif, TTCN_3_GAA (Schindler and Darnell Jr., 1995). We therefore searched for motif enrichment in the differentially bound peaks using STREME (Bailey, 2021) (Figure 4E). This confirmed enrichment of the STAT consensus motif capable of binding canonical STAT5 dimers. Many tools, including STREME, however do not account for one peak containing multiple motifs and since the STAT members are also able to bind to two adjacent motifs as a tetramer (Xu, Sun and Hoey, 1996), we next screened for peaks associated with multiple STAT motifs using RSAT (Thomas-Collier et al., 2011). Dyad enrichment was identified in the differentially bound peaks, thus indicating the possibility that the binding regions could be targeted in WT cells by STAT5 tetramers in addition to the classical dimers (Figure 4E). Finally, we assigned peaks to potential genes targets, using GREAT (McLean et al., 2010). GREAT also allows one peak to be assigned to multiple potential genes if enough evidence supports the annotation. GREAT assigned 136 candidate genes that could be under transcriptional control at the 81 differentially bound sites.

To link differential STAT5 binding to altered gene expression we next performed RNA-sequencing on matched CUT&RUN samples and identified a total of 2,874 significant changes in gene expression, with an FDR <0.01 and fold change >1.5. Of these, 1,588 genes increased and 1,286 decreased in expression in ACK KO cells (Figure 4F). To understand the biological pathways implemented by these gene sets we first used oncoEnrichR (Nakken et al., 2021) to probe the gene set downregulated in ACK KO cells. Significant GO terms affected in ACK KO cells include regulation of cell shape, adhesion, differentiation and protein modification (Figure 4G). These findings were also corroborated using GSEA (Subramanian et al., 2005). 15 Hallmark gene sets in GSEA were enriched in WT cells including IL-2-STAT5 and IL-6-STAT3 signalling, in addition to the GSEA GO Biological Process gene set for the regulation of Rho protein signal transduction (Figure 4H). WT cells are also enriched in GSEA Reactome gene sets for Rac1, Cdc42 and RhoA GTPase cycles, in addition to Rac1 activation (Figure S2B). This analysis confirmed that loss of ACK in K562s affects STAT signalling, as well as disrupting Rho family signalling, as would be expected for a Rho family effector.

Upon examining the overlap of STAT5 CUT&RUN lost peaks and genes downregulated in the RNA-seq data in ACK KO cells, a total of 31 genes were linked to 25 lost STAT5 binding events, which are annotated in the volcano plot in Figure 4F. This gene set constitutes the ACK-STAT5 signature gene set for K562s and is detailed in Table 1. Peaks assigned to downregulated genes are annotated for evidence of the peak being located at a promoter or enhancer location, in addition to the identification of STAT DNA binding motifs and spacings within the peak. These data highlight that the majority of ACK-driven binding events are at sites containing tandem STAT motifs, indicative of sites of potential STAT5 tetramer binding. Full analysis of the CUT&RUN and RNA-sequencing data, including statistical analysis, for this signature set are detailed in supplementary Table 2. This includes motif analysis within peaks where no STAT motifs are found and thus predict alternative transcription factors working in collaboration with STAT5 at this location (Jolma et al., 2013).

**Table 1:**
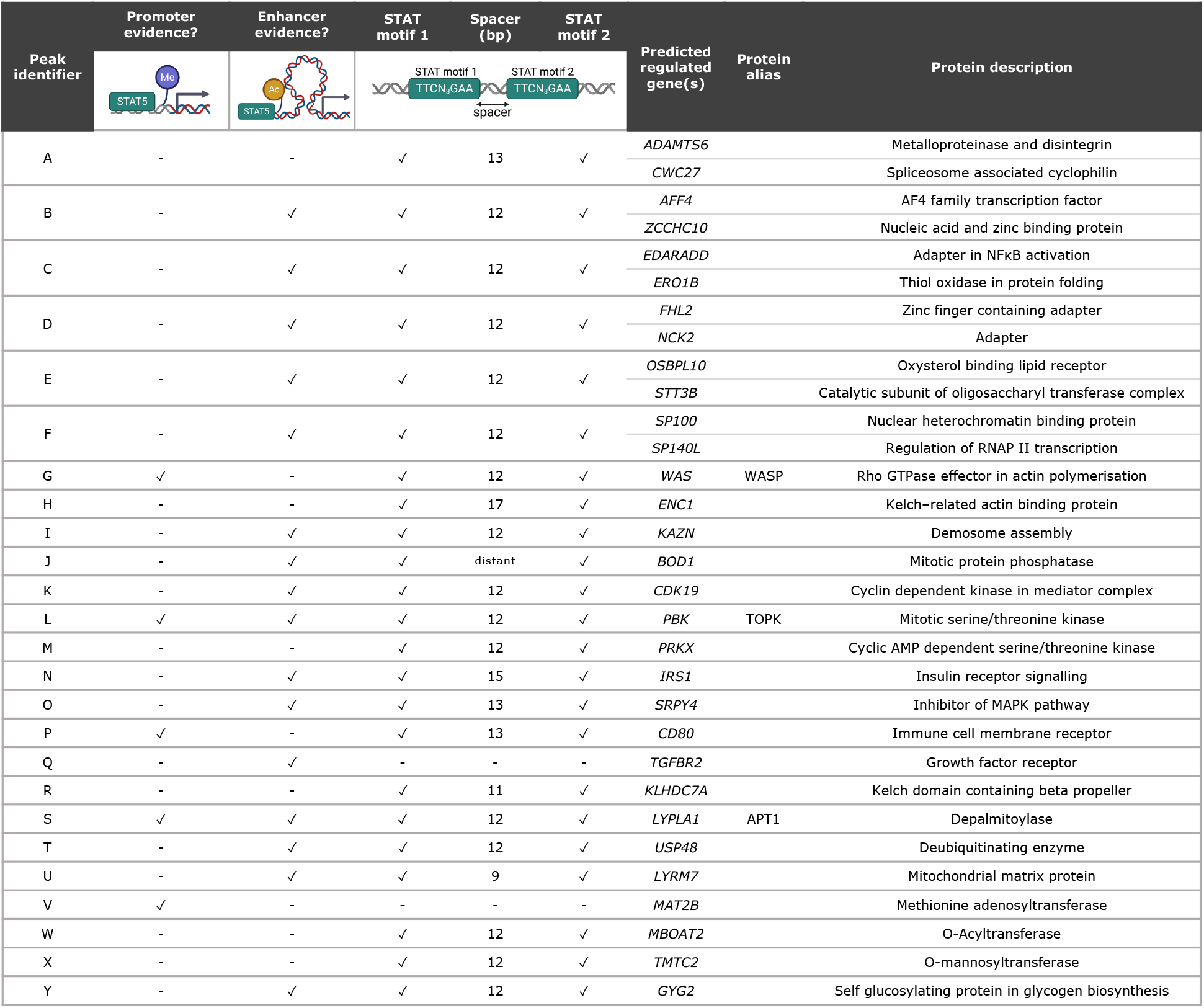
ACK driven STAT5 signature in K562s.

| Peak identifier | Promoter evidence? | Enhancer evidence? | STAT motif 1 | Spacer (bp) | STAT motif 2 | Predicted regulated gene(s) | Protein alias | Protein description |
| --- | --- | --- | --- | --- | --- | --- | --- | --- |
| A | - | - | ✓ | 13 | ✓ | *ADAMTS6* |  | Metalloproteinase and disintegrin |
|  |  |  |  |  |  | *CWC27* |  | Spliceosome associated cyclophilin |
| B | - | ✓ | ✓ | 12 | ✓ | *AFF4* |  | AF4 family transcription factor |
|  |  |  |  |  |  | *ZCCHC10* |  | Nucleic acid and zinc binding protein |
| C | - | ✓ | ✓ | 12 | ✓ | *EDARADD* |  | Adapter in NFκB activation |
|  |  |  |  |  |  | *ERO1B* |  | Thiol oxidase in protein folding |
| D | - | ✓ | ✓ | 12 | ✓ | *FHL2* |  | Zinc finger containing adapter |
|  |  |  |  |  |  | *NCK2* |  | Adapter |
| E | - | ✓ | ✓ | 12 | ✓ | *OSBPL10* |  | Oxysterol binding lipid receptor |
|  |  |  |  |  |  | *STT3B* |  | Catalytic subunit of oligosaccharyl transferase complex |
| F | - | ✓ | ✓ | 12 | ✓ | *SP100* |  | Nuclear heterochromatin binding protein |
|  |  |  |  |  |  | *SP140L* |  | Regulation of RNAP II transcription |
| G | ✓ | - | ✓ | 12 | ✓ | *WAS* | WASP | Rho GTPase effector in actin polymerisation |
| H | - | - | ✓ | 17 | ✓ | *ENC1* |  | Kelch-related actin binding protein |
| I | - | ✓ | ✓ | 12 | ✓ | *KAZN* |  | Demosome assembly |
| J | - | ✓ | ✓ | distant | ✓ | *BOD1* |  | Mitotic protein phosphatase |
| K | - | ✓ | ✓ | 12 | ✓ | *CDK19* |  | Cyclin dependent kinase in mediator complex |
| L | ✓ | ✓ | ✓ | 12 | ✓ | *PBK* | TOPK | Mitotic serine/threonine kinase |
| M | - | - | ✓ | 12 | ✓ | *PRKX* |  | Cyclic AMP dependent serine/threonine kinase |
| N | - | ✓ | ✓ | 15 | ✓ | *IRS1* |  | Insulin receptor signalling |
| O | - | ✓ | ✓ | 13 | ✓ | *SRPY4* |  | Inhibitor of MAPK pathway |
| P | ✓ | - | ✓ | 13 | ✓ | *CD80* |  | Immune cell membrane receptor |
| Q | - | ✓ | - | - | - | *TGFBR2* |  | Growth factor receptor |
| R | - | - | ✓ | 11 | ✓ | *KLHDC7A* |  | Kelch domain containing beta propeller |
| S | ✓ | ✓ | ✓ | 12 | ✓ | *LYPLA1* | APT1 | Depalmitoylase |
| T | - | ✓ | ✓ | 12 | ✓ | *USP48* |  | Deubiquitinating enzyme |
| U | - | ✓ | ✓ | 9 | ✓ | *LYRM7* |  | Mitochondrial matrix protein |
| V | ✓ | - | - | - | - | *MAT2B* |  | Methionine adenosyltransferase |
| W | - | - | ✓ | 12 | ✓ | *MBOAT2* |  | O-Acyltransferase |
| X | - | - | ✓ | 12 | ✓ | *TMTC2* |  | O-mannosyltransferase |
| Y | - | ✓ | ✓ | 12 | ✓ | *GYG2* |  | Self glucosylating protein in glycogen biosynthesis |

We have therefore confirmed that the reduction in pTyr-STAT5 observed in ACK KO K562 lysates results in an altered STAT5 transcriptional landscape with global losses in STAT5-DNA binding, as well as reduced histone methylation and acetylation. Strikingly, the majority of STAT5 binding events lost in ACK KO cells are at putative enhancer locations that could also be potential locations for STAT5 tetramer binding. By combining CUT&RUN data with RNA-sequencing, we have also been able to link lost STAT5 binding events with downregulated gene expression.

### Validation of ACK-driven STAT5 targets

We next sought to corroborate a subset of the ACK-STAT5 signature targets identified in our analysis. Five targets were selected: *WAS, LYPLA1, PBK, SP100* and *AFF4*, based on their high predicted WT protein expression from the RNA sequencing data (Supplementary Table 2) and antibody availability. Genome analysis matched with CUT&RUN data, confirmed a reduction or complete ablation of STAT5 DNA binding in a region linked to each gene and showed that all contained a STAT5 dyad motif spaced by the optimal 12bp (Sathyanarayana et al., 2016) (Figure 5A). All genes were additionally in the vicinity of an active histone mark. RNA-sequencing data for each gene showed that all had significant reductions in mRNA expression in ACK KO cells (Figure 5B). Finally, protein expression levels were confirmed to be reduced in ACK KO cells (Figure 5C), validating their identification as ACK-driven STAT5 transcriptional targets in K562 cells.

**Figure 5:**
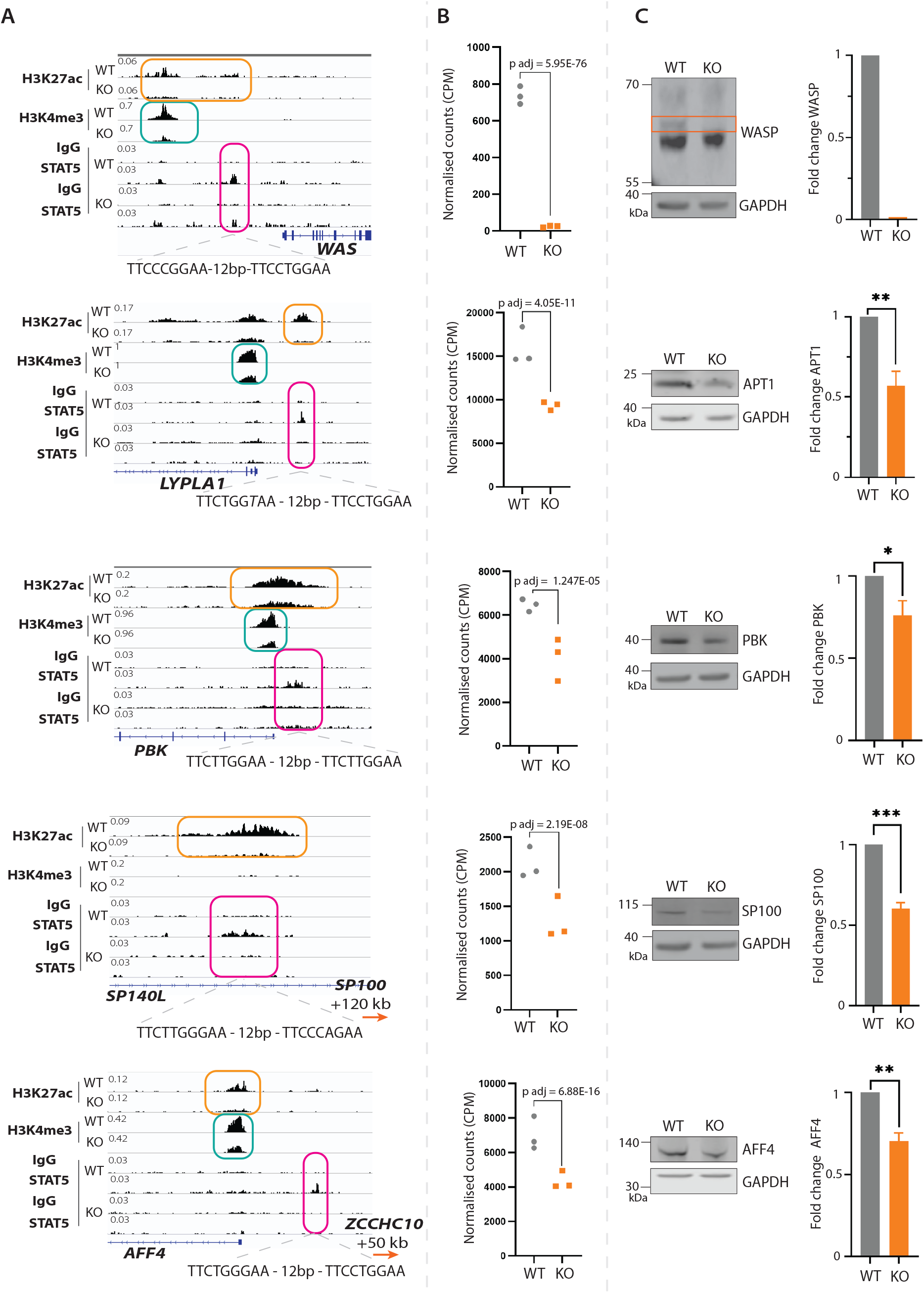
Validation of ACK-driven STAT5 targets in K562s. **(A)** Integrated genomics viewer (IGV) images of STAT5 (pink), H3K4me3 (cyan) and H3K27ac (orange) peaks at loci near to potential gene TSS. The STAT binding motif located within each peak is detailed below. For the peak assigned to *LYPLA1,* one of the motifs has one base divergent from the canonical motif, shown in italics, but this would still allow tetramer binding. **(B)** Graph of normalised counts (CPM) from RNA sequencing of each gene, with statistical significance calculated from DEseq2 and adjusted p value shown. Each point represents data from a biological replicate. **(C)** Protein expression of each target was confirmed by immunoblot and changes compared by densitometry. GAPDH is a loading control. N=3, error bars represent SEM. Statistical significance by unpaired t-test where * p <0.05, ** p <0.01 and *** p <0.001. Letter box (orange) around predicted WASP protein band, protein loss was also confirmed by an additional antibody (not shown).

Many potential phenotypes regulated by ACK-STAT5 were identified in our analysis of the ACK KO K562s, which could contribute to CML. For example, GSEA identified enrichment of the hallmark epithelial mesenchymal transition (EMT) gene set in KO cells (Figure 6A), an important form of cellular remodelling in the transition to a migratory phenotype (Potenta, Zeisberg and Kalluri, 2008). Upregulation of vimentin, a classical EMT marker, was also identified by RNA-sequencing and this was confirmed at the protein level by immunoblot (Figure 6B).

**Figure 6:**
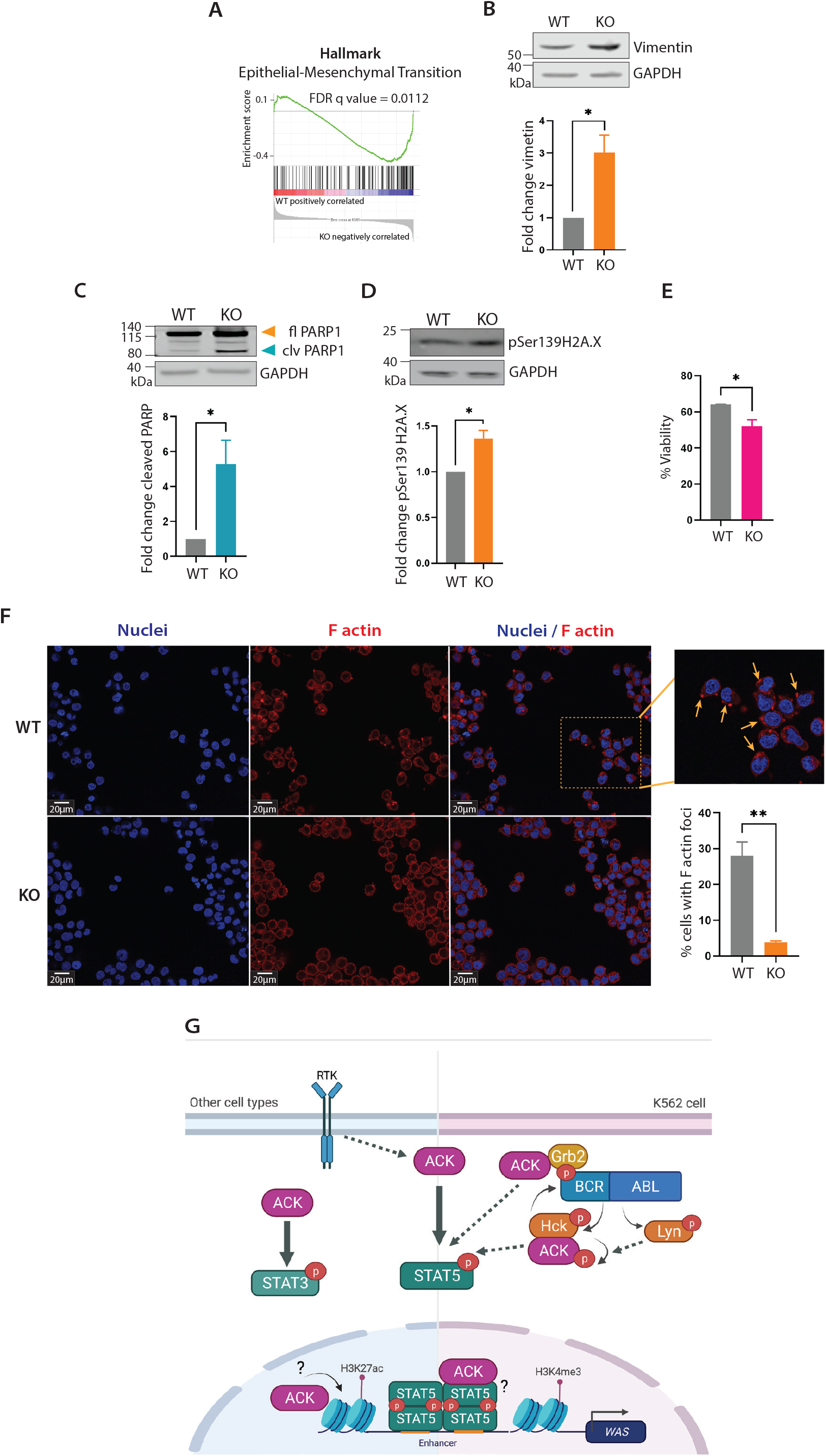
Characterisation of the ACK KO phenotype. **(A)** GSEA plot of Hallmark EMT gene set enrichment in WT K562 cells, showing peak weighting towards the RHS, indicating enrichment in the KO cells. **(B)** Expression levels of vimentin confirmed by immunoblot. GAPDH is a loading control. Quantification to confirm changes in expression by densitometry. N=3, error bars represent SEM. Statistical significance determined by student t-test where * p<0.05. **(C)** Expression levels of full length (fl) and cleaved (clv) PARP1 confirmed by immunoblot. GAPDH is a loading control. Quantification to confirm changes in expression by densitometry. N=3, error bars represent SEM. Statistical significance determined by student t-test where * p<0.05. **(D)** Expression levels of phosphoSer139-H2A.X, confirmed by immunoblot. GAPDH is a loading control. Quantification to confirm changes in expression by densitometry. N=3, error bars represent SEM. Statistical significance determined by student t-test where * p<0.05. **(E)** Percentage viability of cells after 48 hours treatment with 10 μM imatinib mesylate (IM), determined using an automated Beckman cell counter. N=3, error bars represent SEM. Statistical significance by unpaired t-test where * p <0.05. **(F)** Confocal microscopy analysis to monitor changes in F-actin between WT and F1 ACK KO cells. Nuclei were stained with Hoechst (blue) and F-actin with phalloidin (red). Cortical F-actin foci are shown magnified in the inset and identified with arrows. Images acquired using x60 water lens. Scale bars of 20 µM shown. Quantification of cells with the F-actin signal in intense foci was carried out using Columbus image analysis tool. N=3, error bars represent SEM. Statistical significance determined by one-way ANOVA followed by Tukeys multiple testing correction. **** p <0.001. **(G)** Cartoon of possible STAT5 activation routes in K562 cells. ACK contributes to STAT5 phosphorylation in K562 cells, likely acting as the driver in BCR-ABL induced STAT5 activation. BCR-ABL can phosphorylate SFK member Hck. ACK is a known Hck substrate and binding partner and could mediate BCR-ABL-Hck driven activation of STAT5. Hck phosphorylates BCR at Tyr177, a known docking site for adapter Grb2, and additional ACK interacting protein. BCR-ABL can also phosphorylate Lyn, an additional ACK interacting protein. In the nucleus, ACK could direct STAT5 transcription by acting as a transcriptional co-activator in complex with STAT5 or by directing epigenetic changes read by STAT5. ACK can also activate STAT family members in other disease cases and in normal physiological signalling. Figure created in BioRender.

Since genome instability due to defective DNA damage response (DDR) is a classic cancer hallmark, we also investigated whether ACK KO cells have altered DDR pathways (Negrini, Gorgoulis and Halazonetis, 2010). Disruption of pTyr-STAT5 in K562s has also previously been seen to result in the cleavage of poly(ADP-ribose) polymerase 1 (PARP1) (Zhang et al., 2021). We observed increased cleaved PARP1, a nuclear repair enzyme recruited early in the detection of DNA damage (Figure 6C) (Chaudhuri and Nussenzweig, 2017) and increased phosphor-H2A.X (Ser139), a key marker of increased levels of unrepaired double strand breaks (Rogakou et al., 1999) (Figure 6D).

Additionally, we tested the drug sensitivity of ACK KO cells to imatinib mesylate (IM), a common BCR-ABL targeting CML cancer drug, since STAT5 activation levels have been linked to IM responsiveness (Cheng et al., 2018). ACK KO cells were shown to have a modest but significant increase in sensitivity to 10 μM IM treatment (Figure 6E).

ACK KO cells were notably more adhesive when handled and with pathways involving actin organization, cell adhesion and Rho family activation being enriched in WT cells, we next investigated F-actin levels by confocal microscopy. A proportion of WT cells had intense cortical F-actin foci, which have been previously observed in the literature in BCR-ABL expressing cells and suggested to be nascent invadopodia (Figure 6F, expansion) (Li et al., 2007; Daubon et al., 2012). The percentage of cells with intense cortical F-actin foci was significantly reduced in ACK KO cells (Figure 6F lower right). WASP, one of the validated ACK-driven STAT5 targets we analysed (Figure 5), is directly involved in actin polymerization and cytoskeleton organization (Zigmond, 2000) and thus could contribute to this phenotype, but additionally, RNA sequencing has demonstrated ACK loss to have a global effect on the expression of a plethora of proteins critical for Rho family signalling, the master regulators of actin polymerization.

## Discussion

In this study we have shown ACK is a major contributor to oncogenic STAT activation in a CML cell line and thus a potential therapeutic node to dampen a hyperactivated STAT5 system.

In human cells we first demonstrated that ACK interacts with STAT3, STAT5A and STAT5B, resulting in the tyrosine phosphorylation required for activation. We showed that in the presence of ACK, HEK293T cells have increased nuclear pTyr-STAT, resulting in increased transcriptional activity. Previous reports in the literature have shown a biological relationship between ACK and STAT1 in Huh7 liver cells (Fujimoto et al., 2011), in addition to an ACK-STAT1/3 relationship in HEK293T cells that was reliant upon HSP90 (Mahendrarajah et al., 2017). STAT3 was also recently shown to interact with exogenously expressed biotinylated ACK in a triple negative breast cancer cell line (Tahir et al., 2021). Our study builds on these and reports a relationship between ACK and additional STAT family members, STAT5A and STAT5B.

We also demonstrate, to our knowledge, the first endogenous ACK-STAT interactions, specifically in haematological disease cell models. An ACK-STAT3 interaction was seen in HEL and HDLM-2 cell lines, and ACK-STAT5 in K562s. The role of ACK in haematological disease is well studied, with ACK even described as a genetic driver in acute leukaemia (Maxson et al., 2015). HDACi treated acute lymphoblastic leukaemia (ALL) cells carrying an ETV6-RUNX1 fusion have altered ACK expression (Starkova et al., 2007). N-RAS mutated AML and ALL cells are sensitive to the antileukaemic compound GNF7 which acts in this context via specific suppression of the kinase activity of ACK and the serine/threonine kinase GCK (Nonami et al., 2015). Furthermore, juvenile myelomonocytic leukaemia (JMML) cells have been shown to require ACK for survival, with ACK phosphorylating and activating the JMML driver, PTPN11(SHP2) (Jenkins et al., 2018). ACK is also implicated in chronic forms of leukaemia with BCR-ABL negative CML and chronic neutrophilic leukaemia (CNL) cells expressing C-terminal truncated CSF3R showing sensitivity to ACK targeting (Maxson et al., 2013). In K562s, ACK has been identified as a survival gene by pooled CRISPR screening, in addition to being one of the top 20 active kinases in K562s by inferred kinase activity (INKA) analysis (Liu et al., 2019; Beekhof et al., 2019). Thus, ACK is a strong contender for having therapeutic potential in leukaemia.

We have shown that ACK interacts with STAT5 in K562 cells and contributes to the pool of transcriptionally active, nuclear STAT5. We confirmed global loss of STAT5 DNA binding in ACK KO cells, as well as global reduction in histone methylation and acetylation, highlighting the large-scale change in the transcriptional landscape as a consequence of ACK loss. *In silico* analysis of the differences in DNA regions bound by STAT5 between WT and ACK KO cells highlighted that the majority of ACK-driven STAT5 binding events are at potential enhancer binding locations and interestingly at potential STAT5 tetramer binding sites. STAT tetramers form through N-terminal domain interactions and bind cooperatively to adjacent STAT motifs, where the most frequently observed spacer is 12 bp, a spacing seen in 60% of the sites identified in this study (76% with 11-13 bp spacings) (Meyer et al., 1997; John et al., 1999; Sathyanarayana et al., 2016). Very few biological roles have been identified for STAT tetramers, however STAT5 tetramers have a known role in leukaemia development. A leukaemia mouse model no longer develops histological abnormalities on expression of a Δ136-caSTAT5 mutant that can no longer form tetramers, compared to full disease with caSTAT5 (Moriggl et al., 2005). Additionally, a double knock in *Stat5a/Stat5b* tetramer deficient mouse model confirms that STAT5 tetramers drive T-cell proliferation along with natural killer cell maturation and survival (Lin et al., 2012; Lin et al., 2017). It is therefore notable that many of the ACK-driven STAT5 binding events we have identified are at putative tetramer binding sites.

Additionally, we performed RNA-sequencing to identify global changes in gene expression due to loss of ACK. Gene set analysis highlighted that key pathways involved in actin polymerization, cell shape and adhesion were impacted in KO cells. Additionally, signalling from the Rho family of GTPases, of which ACK itself is an effector, is profoundly affected in ACK KO cells. By combining hits from differential STAT5 DNA binding and RNA-sequencing, we identified 31 ACK-driven STAT5 targets in the K562 cell line. We propose this cohort as the ACK-STAT5 signature set in K562 cells. Some of the targets we identified (Table 1) have known links to STAT5, such as CD80, expression of which is regulated by STAT5 in Toll-interacting protein deficient neutrophils, dendritic cells and cutaneous T-cell lymphoma (CTCL), independent of JAK activity (Zhang et al., 2019; Tormo and Gauchat, 2013; Zhang et al., 2014). STAT5 is also known to regulate PBK by direct promoter binding during pregnancy (Cao et al., 2021). Nevertheless, the majority of hits we identified are previously unexplored STAT5 targets. Not only does this signature set indicate pathways that could be interrogated further therapeutically, it also has the potential to be used as a diagnostic for ACK-driven disease

We validated *WAS, LYPLA1, PBK, SP100 and AFF4* as ACK-driven STAT5 targets to the protein level. *WAS* encodes Wiskott-Aldrich syndrome protein (WASP) that, like ACK, is an effector of the Rho family GTPase Cdc42 and regulates actin polymerization. *LYPLA1,* encodes acetyl protein thioeastease 1 (APT1), a depalmitoylase with targets including H-Ras and Gα, and thus regulates the localization and function of key signalling proteins (Camp et al., 1994; Dekker et al., 2010). *PBK* encodes PDZ binding kinase (PBK), a mitotically active serine/threonine kinase (Abe et al., 2000; Gaudet, Branton and Lue, 2000). *SP100* encodes speckled protein 100 which is bound with promyelocytic leukaemia factor (PML) in nuclear structures called PML nuclear bodies (PML-NBs) (Sternsdorf et al., 1999). *AFF4* encodes AF family member 4 (AFF4) which is a scaffold and key component of the super elongation complex (SEC) that regulates RNAP II transcription by releasing transiently paused RNAP II (Lin et al., 2010). Two of these validated hits, *SP100* and *AFF4*, were assigned to peaks that were in the vicinity of another downregulated gene, *SP140L* and *ZCCHC10* respectively. Due to antibody unavailability, we were unable to confirm if they too were downregulated at the protein level. Confirmation of this however would demonstrate that one STAT5 binding event can have dual gene transcriptional control.

Many of the validated hits have known roles in cancer, especially in many types of leukaemia, including, but not exclusively, BCR-ABL driven leukaemia. APT1 is implicated in AML, as well as being described as an oncoprotein in chronic lymphocytic leukaemia (CLL) (Li et al., 2018; Berg et al., 2015). PBK has also been identified as a therapeutic target in FLT3-ITD mutated AML and its expression is altered in BCR-ABL expressing cell lines upon imatinib treatment (Alachkar et al., 2015; Uchida et al., 2019). AFF4 was shown to be a critical part of the SEC in MLL-fusion protein driven leukaemia (Lin et al., 2010). Transcriptional regulation of WASP has also been demonstrated, for example by STAT3 in anaplastic large cell lymphoma (ALCL) and the ETS transcription factor, FLI1 in HELs. Here we add STAT5 in K562s to this list (Menotti et al., 2019; Wang et al, 2021).

Many of our validated hits are also implemented in solid state cancers, suggesting a wider role for this identified signalling axis. For example, APT1 is important for non-small cell lung cancer proliferation, and PBK for *in vivo* colorectal cancer growth, two classes of cancer where ACK is additionally frequently overexpressed (Mohammed et al., 2019; Zhu et al., 2007; Fox, Crafter and Owen, 2019).

We have shown that loss of ACK in K562 cells results in reduced cell proliferation, with the validated hits WASP, AFF4 and PBK also previously linked to K562 proliferation (Weber et al., 2015; Uchida et al., 2019; Liu et al., 2019). ACK KO K562s also have increased expression of EMT markers, increased sensitivity to imatinib treatment, potentially defective DDR and reduced F-actin polymerization.

One of the validated hits, WASP, is a known ACK interacting protein and substrate, subject to both ACK-driven serine and tyrosine phosphorylation (Yokoyama and Miller, 2005). Further integration of the WASP network into ACK signalling was evident in our data: transcription of the WASP interacting protein (WIP), a stabiliser of WASP levels (de la Fuente et al., 2007), was also strongly downregulated in ACK KO cells. Additionally, WASP overexpressing K562 cells show strong upregulation of the protein tyrosine phosphatase PTPRC (Looi et al., 2014) and PTPRC was strongly downregulated in ACK KO K562. WASP KO K562s also have altered PMA-induced F-actin polymerization (Toscano et al., 2013). Therefore the variety of phenotypic changes we have observed in ACK KO cells demonstrate that ACK contributes to numerous signalling pathways critical to multiple stages of cancer progression.

Characteristically K562s express constitutively active BCR-ABL fusion protein which drives constitutive STAT5 phosphorylation (Chai, Nichols and Rothman, 1997; de Groot et al., 1999). The molecular mechanism of BCR-ABL driven STAT5 activation however is not fully understood (Figure 6G). It is believed to be indirect since no direct interaction between STAT5 and BCR-ABL has been observed, suggesting that an intermediatory is required for driving STAT5 activation (Nieboroska-Skorska et al., 1999). The canonical STAT5 activator JAK2, has been ruled out as a mediator since expression of JAK2 kinase dead or dominant negative mutants in CML cells have no effect on pTyr-STAT5 levels, in addition to a JAK2 deletion murine model having no alteration to disease progression (Schafranek et al., 2015; Ilaria and Van Etten, 1996; Xie et al., 2001; Hantschel et al., 2012). It has been postulated that BCR-ABL drives STAT5 activation via the Src family kinases (SFKs), since K562 cells treated with a SFK inhibitor show impeded proliferation and ablated pTyr-STAT5 (Wilson et al., 2002). BCR-ABL is known to activate two SFKs, Hck and Lyn, which are required for BCR-ABL induced STAT5 phosphorylation (Warmuth et al., 1997; Klejman et al.; 2002; Lionberger, Wilson and Smithgal, 2000; Danhauser-Reidl et al., 1996). The kinase activity of Hck is also required for BCR-ABL mediated transformation of DAGM (myeloid leukaemia) cells (Lionberger, Wilson and Smithgal, 2000). Hck can phosphorylate BCR at Tyr177, a known docking site for Grb2, which also interacts with ACK (Warmuth et al.,1997; Satoh et al., 1996). Since ACK is also known to be a direct Hck substrate (Yokoyama and Miller, 2003), it is credible that ACK is the missing link in BCR-ABL dependent STAT5 activation via the SFKs, with hyperactive BCR-ABL in CML providing the ACK activatory signal (Figure 6G). ACK could also be able to activate other STAT family members in other disease scenarios and indeed in normal physiological signalling, since ACK lies downstream of other activators such as receptor tyrosine kinases (RTK) (Figure 6G).

One obvious question remains; how does ACK influence STAT5 binding? ACK is known to influence epigenetic changes such as histone modifications that result in the recruitment of transcriptional complexes (Mahajan et al., 2017). With clear losses in global histone methylation and acetylation in ACK KO K562s, the chromatin landscape in these cells is clearly widely altered. Does ACK influence changes in STAT5 binding by stimulating epigenetic change at histones? (Figure 6G). In castration resistant prostate cancer (CRPC) cell models where ACK deposits pTyr88 Histone 4 marker sites, STAT5 was identified in *de novo* motif searching at these locations, as a potential transcription factor recruited to these new locations (Mahajan et al., 2017). Potentially ACK could be altering chromatin marking to allow accessibility for STAT5 in K562 cells too. Conversely, ACK is known to influence the AR transcriptional signature in prostate cancer models by acting as a direct partner and component of the AR DNA binding complex (Mahajan et al., 2007)(Figure 6G). Thus, similarly, ACK could be part of the chromatin associated STAT5 complex at loci in CML, directing STAT5 to its signature set of targets.

In summary, we have shown that ACK can interact, stimulate tyrosine phosphorylation, nuclear translocation and transcriptional activation of STAT3, STAT5A and STAT5B in HEK293T cells. In the K562 cell line we have also demonstrated that ACK interacts with STAT5 and drives a pool of nuclear, transcriptionally active STAT5. We have identified 31 ACK-driven STAT5 transcriptional targets, many of which are driven by STAT5 enhancer binding at potential tetramer binding locations and constitute a signature for ACK-STAT5 activity in K562 cells. Thus, we have identified ACK as a driver of oncogenic STAT activation in a CML cell model and thus a potential therapeutic node to dampen hyperactivated STAT5.

## Acknowledgements

We are grateful to Ulrike Künzel, Millie Fox, Sinead Knight, David Fisher, Ian Barrett and Hannah Webb (AstraZeneca, Cambridge UK), in addition to Julia Lindgren and Graham Belfield (AstraZeneca, Gothenburg, Sweden) for their insightful discussions and practical help. We also thank Lawrence Welch for help with paper editing.

---

**Figure S1:**
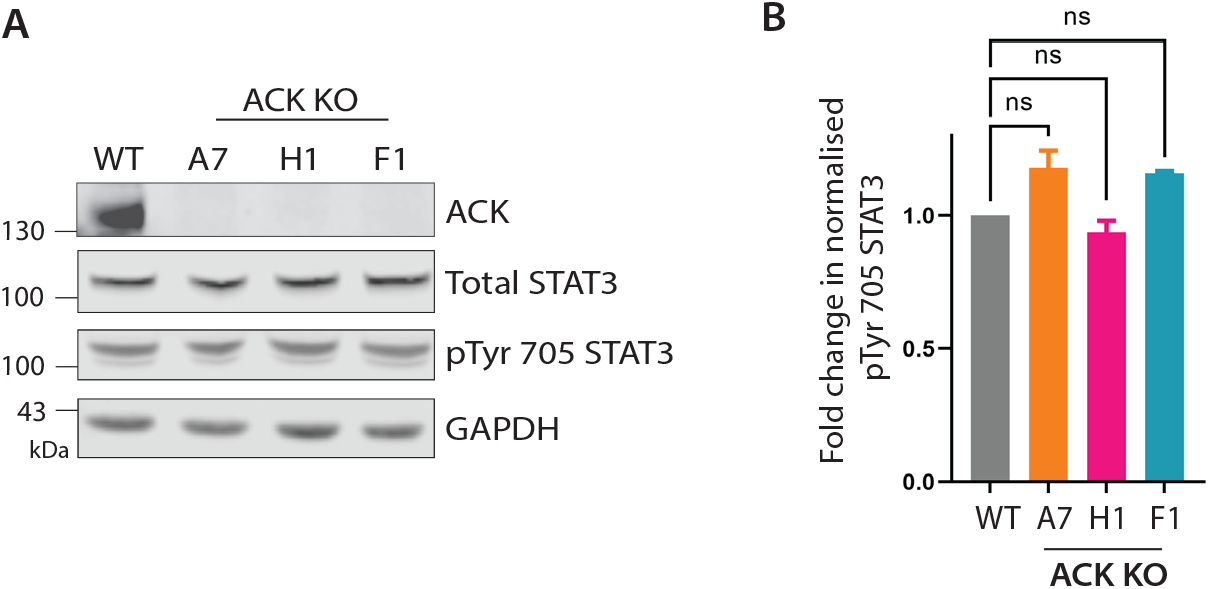
ACK-STAT3 relationship in ACK KO K562s. **(A)** Immunoblot to confirm levels of total STAT3 and pTyr705 STAT3 in WT and ACK KO K562 cell lysates. GAPDH is a loading control. **(B)** Quantification of pTyr705 STAT3 signal normalised to total STAT3 levels by densitometry. N=3, error bar represents SEM. Statistical significance determined by one-way ANOVA followed by Dunnett’s multiple correction test. ns=not significant.

**Figure S2:**
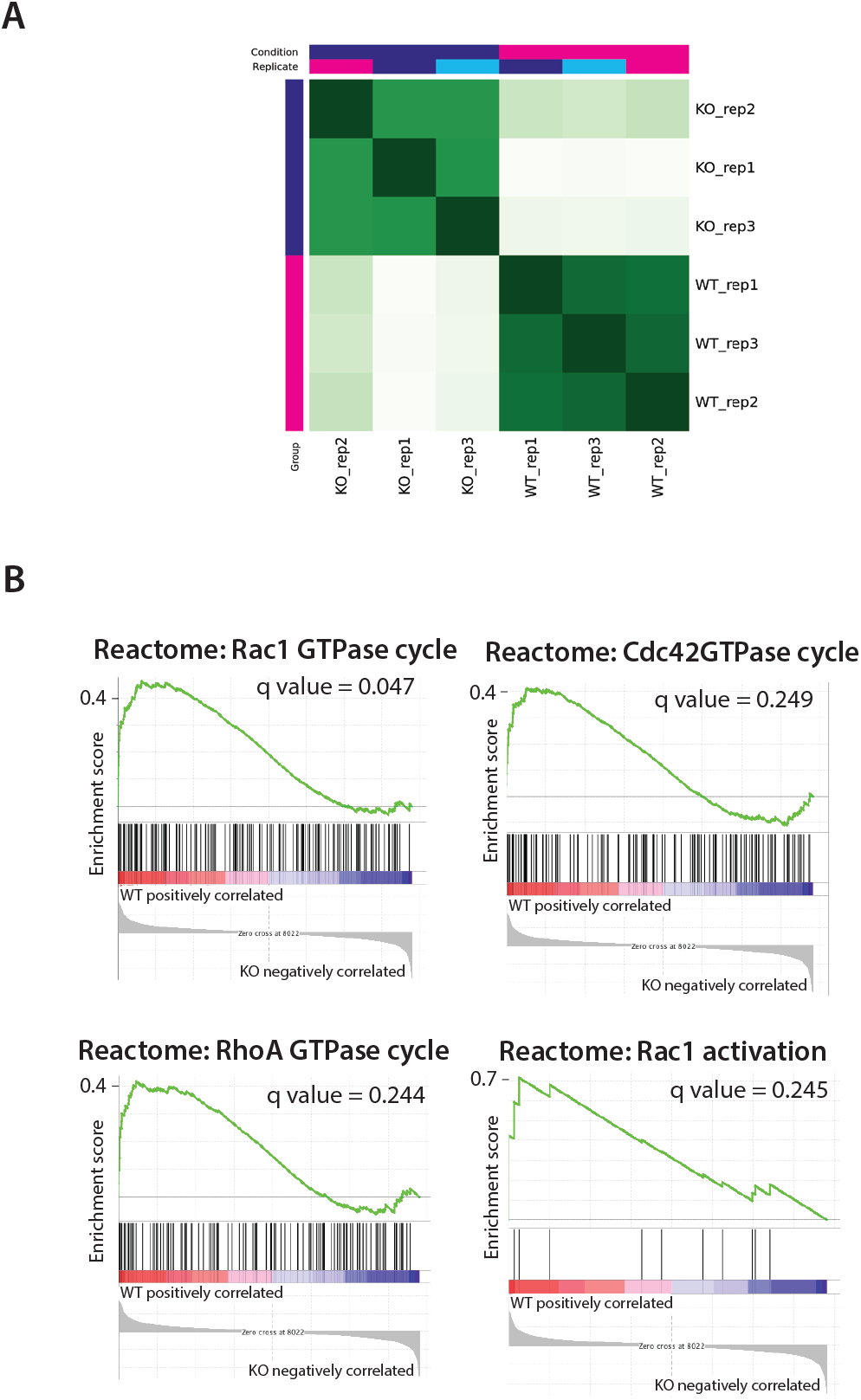
Extended bioinformatic analysis of ACK KO. **(A)** Hierarchical clustering correlation heatmap of significantly differentially bound sites between WT and KO conditions, reflecting the agreement between biological replicates. Two conditions are represented (purple, KO and pink, WT) with 3 biological replicates (replicate 1 purple, replicate 2 pink and replicate 3 blue). Darker shades of green are reflective of higher correlation between biological replicates. All outputs directly from Diffbind. **(B)** Further GSEA plots showing enrichment of Rac1/Cdc42/Rho A GTPase cycles and Rac1 activation in WT K562 cells.

**Supplementary Table 1:**
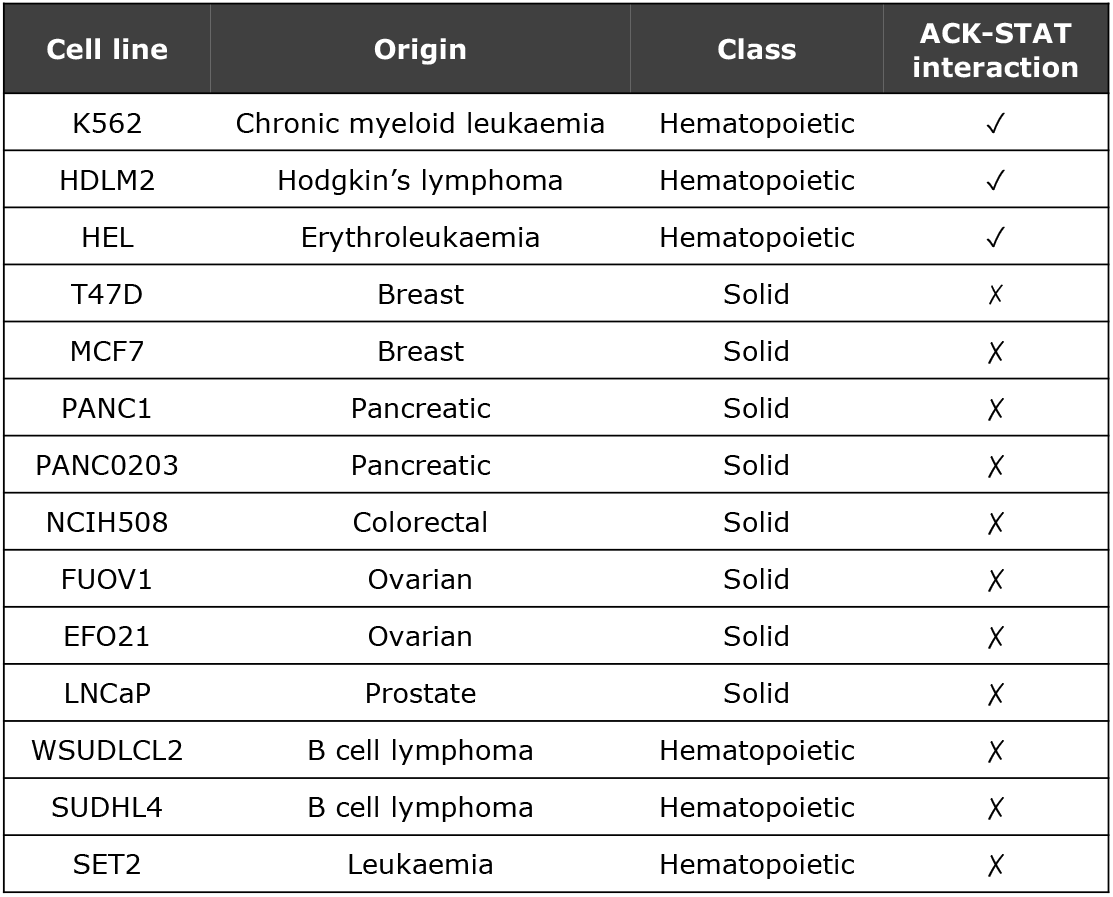
Cell lines tested for endogenous ACK-STAT3 or STAT5 interaction

**Supplementary Table 2:**
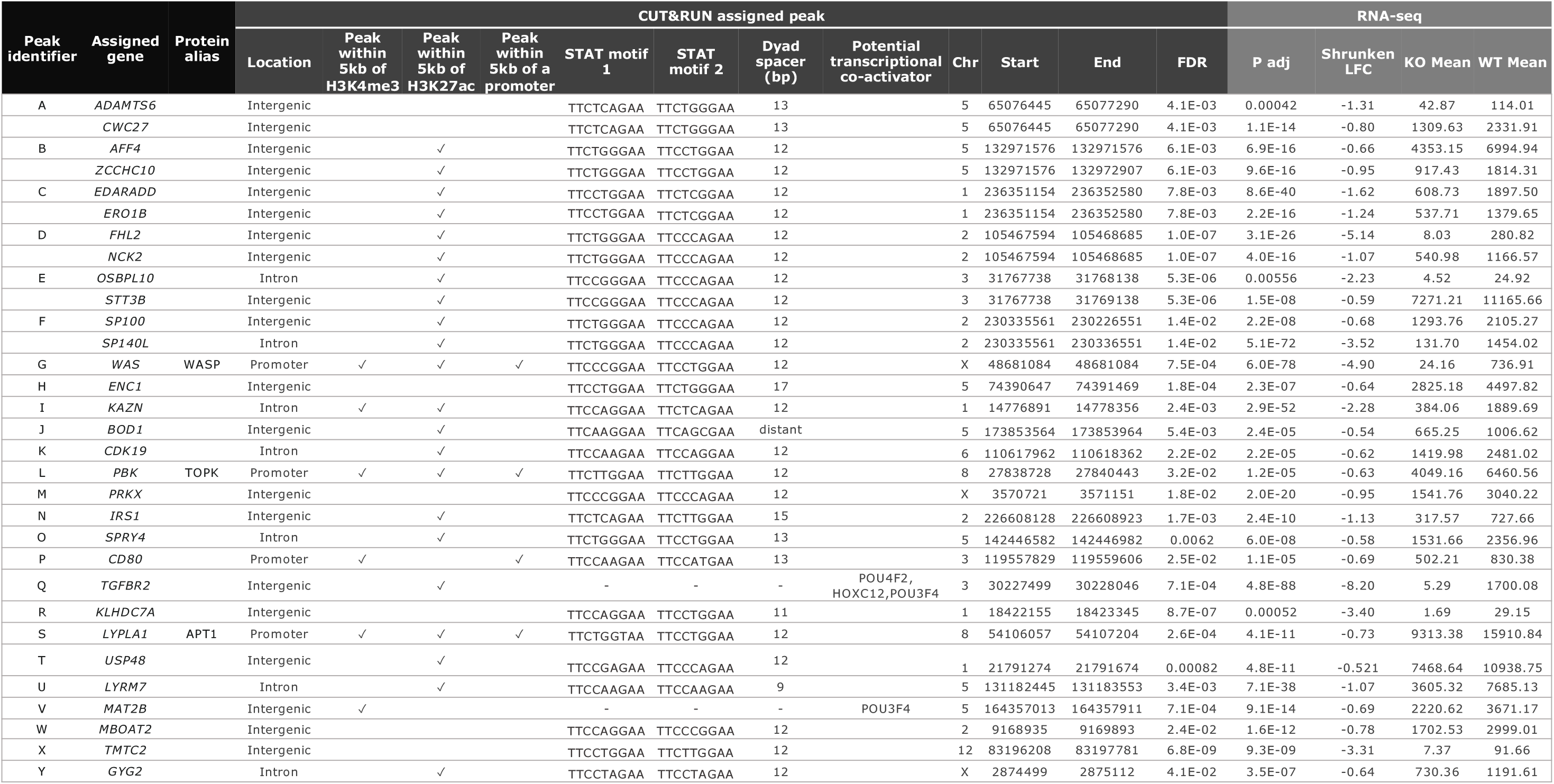
Full detail of the ACK-STAT5 K562 signature.

Supplementary data file 1: DiffBind file from CUT&RUN analysis.

Supplementary data file 2: DESeq2 file from RNA-sequencing.

## Methods

### Cells and reagents

HEK293T, HEK239T/17, HEL, HDLM-2 and K562 cells were obtained from the AstraZeneca cell bank (Alderley Park, UK) and maintained at 37°C in 5% CO_2_ and grown in DMEM or RPMI-40 (Gibco) with 10% fetal bovine serum (ThermoFisher), 2 mM L-Glutamine and 1% Antibiotic-Antimycotic (Gibco). Cells were routinely tested for mycoplasma (Minerva Biolabs). AIM-100 was purchased from Tocris and DMSO from Sigma Aldrich.

Cells were either transfected with PEI (1 μg/3 μg DNA) or lipofectamine 2000 (ThermoFisher) diluted in OptiMEM (Gibco) and incubated for 40 hours before analysis.

### Vectors

pXJ-HA-GFP, pXJ-HA-ACK and pXJ-HA-dACK were gifts from Professor Ed Manser (ICMB, Singapore), pCMV-Renilla from Dr Marc de la Roche (University of Cambridge, UK), pRK5-STAT5A-FLAG and pRK-STAT5B-FLAG from Professor James Irle (St Jude’s Children’s Research Hospital, Memphis, Tennessee) and pcDNA3.1/V5-His-STAT3 TOPO was gifted from Dr Jonathon Wilde, GSK, Stevenage, UK. STAT3 was then subcloned into a pENTR/D-TOPO vector and transferred into a pDEST12.2-FLAG destination vector by Gateway recombination.

### Confocal microscopy

Cells were seeded at a density of 10,000 cells in 96 well plates. HEK239T/17 cells were transfected with Lipofectamine 2000 24 hours post seeding and fixed 40 hours post transfection. K562 cells were fixed 24 hours post seeding. Cells were fixed with a final concentration of 4% paraformaldehyde and blocked with modified blocking buffer (PBS, 1.1% BSA and 0.1% Triton-X). Primary antibodies were incubated overnight diluted in modified blocking buffer. Cells were then washed with phosphate buffered saline-0.1%Tween (PBST) and incubated with secondary antibody and Hoechst for 1 hour prior to imaging at room temperature in the dark. Cells were then re-washed in PBST. Cells were imaged on a Cell Voyager 7000 (Yokogawa) using the x60 water immersion lens and images subjected to quantitative analysis using Columbus Image Analysis software (Perkin Elmer). Intensity thresholds for each antibody were established and cells counted that were below or over determined thresholds.

### Antibodies

The following antibodies were used for western blot: STAT3 (Cell Signalling, CS9139, 1:1000 dilution), pTyr705 STAT3 (Cell Signalling CS9131, 1:2500 dilution), STAT5 (SantaCruz, sc-74442, 1:2500 dilution), pTyr694/699 STAT5 (Cell Signalling, CS9351, 1:5000 dilution), ACK (SantaCruz, SC-28336, 1:1000 dilution), pTyr284 ACK (Merck, 09-142, 1:2500 dilution), GAPDH (Santacruz, SC-47724, 1:1250 dilution), Histone 3 (Abcam, AB1791, 1:50,000 dilution), FLAG-HRP (Sigma Aldrich, A8592, 1:5000 dilution), WASP (SantaCruz, SC-13139, 1:250 dilution), WASP (NovusBio, MAB30701, 1:500 dilution), LYPLA1 (Abcam, AB91606, 1:5000 dilution), PBK (Cell signalling, CS4942, 1:5000) dilution, AFF4 (Proteintech, 14662-1AP, 1:3000 dilution), SP100 (Proteintech, 11377-1AP, 1:400 dilution), PARP1 (Proteintech, 13371-1-AP, 1:1000 dilution), phospho-H2A.X(pSer139) (Cell signalling, CS9718, 1;1000 dilution), vimentin (Cell signalling, CS5741, 1:1000 dilution, goat anti mouse IgG-HRP (Newmarket Scientific, GtxMu-0030DHRPX, 1:5000 dilution), goat anti rabbit IgG-HRP (Newmarket Scientific, GtxRb-0030DHRPX, 1:5000 dilution), goat anti mouse IRDye 800CW (Licor, 926-32210, 1:5000 dilution) and goat anti rabbit IRDye 800CW (Licor, 926-32211, 1:5000 dilution).

The following antibodies were used for immunoprecipitation: FLAG-M2 (Sigma Aldrich, F31651, 1 μg), Mouse IgG (Santacruz, SC-2025, 1 μg) and ACK (Santacruz, SC-28336, 1 μg).

The following antibodies were used for confocal microscopy: ACK (abcam, ab185726, 1:500 dilution), pTyr705-STAT3 (Cell signalling, CS9139, 1:100 dilution), pTyr694/699-STAT5 (BD Bioscience, 61194, 1:100 dilution), Hoescht 3342 (Invitrogen, H3570, 1:40,000 dilution), Phalloidin-488 (Invitrogen, A12379, 1:40 dilution), goat anti rabbit IgG (H+L) cross absorbed Alexa Fluor 488 (Invitrogen, 11008, 1:5000 dilution) and goat anti mouse IgG (H+L) cross absorbed Alexa Fluor 555 (Invitrogen, 21424, 1:5000 dilution).

### Cell proliferation

Cells were seeded at a density of 0.4 × 10^6^ cells / mL and incubated at 37°C in 5% CO_2_ over a period of up to 72 hours. Cell numbers were counted using the Vi-Cell automated cell counter (Beckman) and proliferation calculated as fold change compared to the seeding density.

### Immunoprecipitation

Cells were either lysed in ice-cold lysis buffer (50 mM Tris-HCl pH7.5, 150 mM NaCl, 1 mM EDTA, 1 mM sodium orthovanadate, 1 mM β-glycerophosphate disodium salt hydrate, 1x mammalian PIC (Sigma Aldrich) and 1% Triton-X) or RIPA lysis buffer (ThermoFisher). Lysates were centrifuged at 13,000 × g for 20 minutes at 4°C and supernatant collected. Lysates were pre-cleared by rotating in 20 µL of Protein G Dynabeads (Thermo Scientific) at 4°C for 30 minutes. Lysates were then incubated with either 1 µg FLAG-M2 antibody for 1 hour at 4°C, or appropriate primary antibody overnight. Lysates were then rotated with 30 µL Protein G Dynabeads (Thermo Scientific) at 4°C for 30 minutes before 3X lysis buffer washes. Proteins were then eluted in 30 µL 2x LDS reducing sample buffer and 30 µL PBST. Proteins were analysed by immunoblot.

### Subcellular fractionation

HEK293T cells were collected in 5 mL chilled PBS and pelleted at 1000 × g for 15 minutes. The pellet was resuspended in 0.5 mL ice cold hypotonic buffer (20 mM Tris-HCl pH7.4, 10mM NaCl, 3mM MgCl_2_) and incubated on ice for 15 minutes. 25 µL 10% NP-40 was added and cells vortexed for 10 s. Cells were centrifuged at 1000 × g for 20 minutes and supernatant collected as the cytoplasmic fraction. The pellet was resuspended in 500 µL ice cold hypotonic buffer and re-centrifuged at 1000 × g for 20 minutes. The pellet was resuspended in 500 µL ice cold cell extraction buffer (100 mM Tris-HCl pH 7.4, 100 mM NaCl, 1 mM EDTA, 1 mM EGTA, 0.1% SDS, 1 mM NaF, 2 mM Na_3_VO_4_, 1% Triton-X, 10% glycerol, 0.5% sodium deoxycholate, 20 mM Na_4_P_2_O_7_, 1 mM PMSF, 1x protease inhibitor cocktail (Sigma Aldrich). Samples were kept on ice for 40 minutes, vortexing at maximum speed every 10 minutes. After a final 13,000 × g centrifuge spin for 20 minutes at 4°C, the supernatant was collected as the nuclear fraction. 2x LDS reducing sample buffer was added 1:1 to both cytoplasmic and nuclear fractions and proteins analysed by immunoblot. K562 cells were fractionated using the Subcellular Protein Fractionation Kit for Cultured Cells (ThermoFisher) according to maunfacturer’s instructions.

### Immunoblot analysis

Proteins were separated on 8% SDS-PAGE or 4-12% NuPAGE gels (ThermoFisher) and proteins transferred to a methanol activated PVDF membrane (Immobilon) using a semi dry blotting system (Hoefer Scientific), or an iBlot2 (ThermoFisher). Membranes were blocked in 5% milk in PBST. Protein bands were visualised by enhanced chemiluminescence or by fluorescence using the Odyssey-Cxl (Licor). Densitometry was performed using ImageStudio.

### Luciferase reporter assay

3 × 10^5^ HEK293T cells were seeded in 6 well plates. 24 hours later cells were transiently transfected with HA-ACK (1 µg for STAT3, 0.5 µg for STAT5A and 0.75 µg for STAT5B), 1 µg FLAG-STAT, either 2 µg pGL4.47[luc2P/SIE/Hygro] (Promega) or pGL4.52[luc2P/STAT5RE/Hygro] (Promega) and 20 ng pCMV-Renilla (Promega) using PEI. 24 hours later media was replenished with 0.05% serum media and cells treated with 10 µM AIM-100 (for STAT3), and 1 µM (for STAT5A/B) or DMSO control. 16 hours later cells were lysed using the Dual Luciferase reporter assay system (Promega). Luciferase activity, detected on an Pherastar plate reader (BMG LabTech), was normalized to *Renilla* luciferase activity and transcription levels calculated as fold changes from baseline levels. Lysate samples were also analysed by immunoblot.

### ACK KO K562 cell line generation

CRISPR-Cas9 technology was used to knockout *TNK2* by introducing frameshifts within exon 6 with two guides identified from the V3 Yusa library (guide 1: 5’CGCGACCTGGTCAAGATCG 3’ and guide 2: 5’ GACGACCATTACGTCATGC 3’), and purchased from ThermoFisher. An RNP complex of gRNA, Cas9 (produced by AstraZeneca Protein Science) and tracrRNA (IDT, 1072532) was formed in duplex buffer (IDT, 11010301) and electroporated into 2 × 10^5^ wild type K562 cells using the Neon electroporation system (ThermoFisher) at 1450 V, 10 ms for 3 pulses. 1 week post electroporation, cells were single cell sorted using the CellenONE (Scienion) into 96 well plates in 20% conditioned media, and single cell clones were grown up over a period of weeks. Clones were identified by genotyping PCR (forward primer = 5’ CTCCAGCTCATTCCTCCTGC 3’ and reverse primer = 5’ GGACACAGCATCTATCCCCG 3’) with KO efficiency determined using the TIDE analysis tool (Brinkman et al., 2014). Candidate KO clones were then grown up and whole cell lysates accessed by immunoblot to confirm absence of target protein.

### Cleavage under targets and release using nuclease (CUT&RUN)

Chromatin profiling for WT and ACK KO K562 cells was performed using CUTANA™ ChIC/CUT&RUN Kit (EpiCypher) according to the manufacturer’s protocol with minor modifications. Per replicate, 2 million WT and F1 ACK KO K562 cells were crosslinked in a final concentration of 1% formaldehyde for 1 minute at room temperature before quenching with 125 mM glycine. Samples were cryopreserved and stored at −80°C before use. On the day of processing, Wash Buffer, Cell Permeabilization Buffer and Antibody Buffer were supplemented with 1% Triton X-100 and 0.05% SDS, and used in the following steps. Vials with 2 million cryopreserved cells per sample were placed in a 37°C water bath for slow thawing and washed twice with 400 µL of Wash Buffer. Washed cells were resuspended in 410 µL of Wash Buffer and 100 µL of this solution was aliquoted into four reaction tubes i.e. 5 × 10^7^ cells per CUT&RUN reaction. 100 µL of pre-activated ConA bead slurry was added to each reaction tube and the mixture incubated for 10 min at room temperature to allow cell-bead binding. Cell-bead solution was then buffer exchanged into Antibody Buffer and target-specific antibodies added to individual reaction tubes in the following amounts: 0.5 µg of anti-H3K4me3 (EpiCypher, 13-0041), 1 µg of anti-H3K27ac (EpiCypher, 13-0045), 0.6 µg of anti-Stat5 (Santa Cruz Biotechnology, sc-271542) and 0.5 µg of IgG (EpiCypher, 13-0042) as a negative control. Antibody binding was performed at 4°C overnight with gentle mixing on a nutator. Binding of pAG-MNase, targeted chromatin digestion and DNA release were completed according to the manufacturer’s protocol. To reverse cross-links, CUT&RUN released DNA was supplemented with 0.8 µL of 10% SDS (Merck) and 1 µL of 20 µg/ul Proteinase K (Ambion), and incubated overnight at 55°C using a thermocycler. DNA purification was performed using DNA Cleanup Columns.

Purified, CUT&RUN-enriched DNA was adapted for sequencing using KAPA HyperPrep Kit (Roche) and multiplexed using 1.5 µM custom TruSeq UDI-UMI adapters (Integrated DNA Technologies) with minor modifications to the manufacturer’s protocol. Briefly, 1x bead-based cleanup was followed by an adapter-ligation step and the library was amplified using 14 PCR cycles with a hybrid 10 s anneal/extension parameter at 60°C, to limit amplification of higher molecular weight species. Resulting libraries were quantified on a Fragment Analyzer 5300 system (Agilent), pooled equimolarly to a final 1.9 nM concentration, and sequenced using a 51 bp paired-end setting on a v1.5 SP NovaSeq6000 flowcell (Illumina).

Adaptor sequences were trimmed and poor-quality reads were removed from the ends of reads using Trimmomatic (v0.39) (PMID: 24695404). Additionally, kseq (v20190822) was run to remove additional barcode sequences potentially missed through Trimmomatic processing (PMID: 31500663). Trimmed reads were aligned to the human genome (GRCH38, Gencode v 31) using Bowtie2 (v 2.4.2) with the parameters “--very-sensitive-local” to assign multi-mapped reads to their best alignment and “--dovetail” to permit the alignment of mate pairs that extend past one another (PMID: 22388286). Reads were also aligned to the spike-in *E. coli* genome (K12-MG1655) using Bowtie2 with the additional parameters “--no-overlap” and “--no-dovetail”. Aligned reads were filtered using SAMtools (v0.8.1) (PMID: 19505943) for a minimum quality score of 0. Spike-in normalization was performed using the procedure outlined in Skene et al. 2017 (PMID: 28079019). Briefly, a scale factor was calculated for each sample based on the depth of the corresponding spike-in and Bedtools “genomecov” (v2.30.0) was used to scale target bam files and output a bedgraph file (PMID: 20110278). For genome browser visualization, ucsc “bedGraphToBigWig” (v377) was used (PMID: 20639541). For peak calling, macs2 “callpeak” (v2.2.7.1) and overlapping peaks across samples were merged using HOMER “mergePeaks” (v4.11) (PMID: 18798982, PMID: 20513432). Raw read quality was assessed using FastQC (version 0.11.9) (https://www.bioinformatics.babraham.ac.uk/projects/fastqc/), alignment metrics was assessed using picard (v2.25.0) (https://gatk.broadinstitute.org/hc/en-us/articles/360040507751-CollectAlignmentSummaryMetrics-Picard-) and SAMtools (v1.10), and binding enrichment quality metrics were generated through preseq (v3.1.2) (http://smithlabresearch.org/software/preseq/), deeptools (v3.5.0) (PMID: 27079975) and Phantompeakqualtools (v1.2.2) (PMID: 22955991). Finally, quality metrics were summarized using MultiQC (v1.10.1) (PMID: 27312411). Data processing was managed using Snakemake (v5.26.1) (PMID: 34035898).

Peak annotation was performed using ChIPseeker (v1.30.3) (PMID: 25765347) and differentially bound peaks identified using DiffBind (v3.4.3) (PMID: 22217937). *In silico* motif analysis was performed using STREME (v5.4.1) (PMID: PMID: 33760053) and RSAT (PMID: 22156162). Some analysis was performed on the galaxy web platform, using the public server usegalaxy.org. Overlap of differentially bound peaks with ENCODE K562 identified cCREs was performed online via ENCODE SCREEN registry v3 (screen.encodeproject.org) (PMID: 32728249).

### RNA-sequencing

An additional 1 × 10^6^ WT and ACK KO K562 cells was harvested for each CUT&RUN replicate before crosslinking and used for matched mRNA sequencing. Briefly, total RNA was extracted using Agencourt RNAdvance Tissue kit including DnaseI (Ambion) treatment for 15 min at 37°C on an automated Biomek i7 Hybrid system (Beckman Coulter). RNA quantification and quality control was performed on Fragment Analyzer 5300 system and showed an RNA integrity number (RIN) of 10 for all the samples. mRNA-seq library preparation was performed using KAPA Stranded mRNA-Seq Kit, with KAPA mRNA Capture Beads (Roche) following manufacturer’s protocol with 500 ng of input RNA, 5 min fragmentation time at 95°C, 10 PCR cycles, and multiplexed using TruSeq UD-UMI adapters. Resulting libraries were pooled equimolarly to a final concentration of 1.9 nM. 51 bp paired-end sequencing was performed on a v1.5 SP NovaSeq6000 kit (Illumina).

Libraries were accessed using FastQC (v0.11.7), Qualimap (v2.2.2c) (PMID: 26428292) and SAMtools stats (v1.9). Alignment was performed using STAR (version 2.7.2b) (PMID: 23104886) with alignment against the human genome (GRCh38, Ensembl v100). Sequencing Quality control metrics were obtained using Qualimap (v2.2.2c) (PMID: 26428292) and summarized using MultiQC (v1.9). Trimming of adapters was performed using NGmerge (v0.3) (PMID: 30572828).

A human transcriptome index consisting of cDNA and ncRNA entries from Ensembl v100) was generated and gene abundances were obtained using Salmon (v1.1.0) (PMID: 28263959). The bioinformatics workflow was organized using Nextflow workflow management system (v20.10) (PMID: 28398311) and Bioconda software management tool (PMID: 29967506). For differential gene expression analysis, DESeq2 (v1.26.0) was used with “ashr” (PMID: 27756721) for fold change shrinkage, all in R (v3.6.1) (PMID: 25516281). Gene set enrichment analysis was performed with GSEA (v4.2.0) (PMID:16199517) and OncoEnrichR Cancer-dedicated gene set interpretation (Galaxy v1.0.7 https://oncotools.elixir.no/) (arXiv:2107.13247v2).

## References

1. Abe Y, Matsumoto S, Kito K & Ueda N (2000) Cloning and expression of a novel MAPKK-like protein kinase, lymphokine-activated killer T-cell-originated protein kinase, specifically expressed in the testis and activated lymphoid cells. J. Biol. Chem. 275: 21525–21531

2. Alachkar H, Mutonga M, Malnassy G, Park J-H, Fulton N, Woods A, Meng L, Kline J, Raca G, Odenike O, Takamatsu N, Miyamoto T, Matsuo Y, Stock W & Nakamura Y (2015) T-LAK cell-originated protein kinase presents a novel therapeutic target in FLT3-ITD mutated acute myeloid leukemia. Oncotarget; 6 (32) 33410–33425.

3. Bailey TL (2021) STREME: accurate and versatile sequence motif discovery. Bioinformatics 37: 2834–2840

4. Beekhof R, van Alphen C, Henneman AA, Knol JC, Pham T V, Rolfs F, Labots M, Henneberry E, Le Large TY, de Haas RR, Piersma SR, Vurchio V, Bertotti A, Trusolino L, Verheul HM & Jimenez CR (2019) INKA, an integrative data analysis pipeline for phosphoproteomic inference of active kinases. Mol. Syst. Biol. 15: e8250–e8250

5. Berg V, Rusch M, Vartak N, Jüngst C, Schauss A, Waldmann H, Hedberg C, Pallasch CP, Bastiaens PIH, Hallek M, Wendtner C-M & Frenzel LP (2015) miRs-138 and -424 control palmitoylation-dependent CD95-mediated cell death by targeting acyl protein thioesterases 1 and 2 in CLL. Blood 125: 2948–2957

6. Brinkman, E. K., Chen, T., Amendola, M., & van Steensel, B. (2014). Easy quantitative assessment of genome editing by sequence trace decomposition. Nucleic acids research, 42(22), e168.

7. Buettner R, Mora LB & Jove R (2002) Activated STAT signaling in human tumors provides novel molecular targets for therapeutic intervention. Clin. Cancer Res. 8: 945–954

8. Camp LA, Verkruyse LA, Afendis SJ, Slaughter CA & Hofmann SL (1994) Molecular cloning and expression of palmitoyl-protein thioesterase. J. Biol. Chem. 269: 23212–23219

9. Cao Y, Feng Z, He X et al (2021) Prolactin-regulated Pbk is involved in pregnancy-induced β-cell proliferation in mice. J Endocrinol. 252: 107–123

10. Chai SK, Nichols GL & Rothman P (1997) Constitutive activation of JAKs and STATs in BCR-Abl-expressing cell lines and peripheral blood cells derived from leukemic patients. J. Immunol. 159: 4720 LP –4728

11. Chaudhuri A and Nussenzweig A (2017) The multifaceted roles of PARP1 in DNA repair and chromatin remodelling. Nat. Rev. Mol. Cell Biol. 18: 610–621.

12. Cheng Y, Hao Y, Zhang A, Hu C, Jiang X, Wu Q & Xu X (2018) Persistent STAT5-mediated ROS production and involvement of aberrant p53 apoptotic signaling in the resistance of chronic myeloid leukemia to imatinib. Int. J. Mol. Med. 41: 455–463

13. Clayton NS, Fox M, Vicenté-Garcia JJ, Schroeder CM, Littlewood TD, Wilde JI, Krishnan K, Brown MJB, Crafter C, Mott HR & Owen D (2022) Assembly of nuclear dimers of PI3K regulatory subunits is regulated by the Cdc42-activated tyrosine kinase ACK. J. Biol. Chem. Adance online publication.

14. Corry J, Mott HR & Owen D (2020) Activation of STAT transcription factors by the Rho-family GTPases. Biochem. Soc. Trans. 48: 2213–2227

15. Danhauser-Riedl S, Warmuth M, Druker BJ, Emmerich B & Hallek M (1996) Activation of Src kinases p53/56(lyn) and p59(hck) by p210(bcr/abl) in myeloid cells. Cancer Res. 56: 3589–3596

16. Daubon T, Rochelle T, Bourmeyster N & Génot E (2012) Invadopodia and rolling-type motility are specific features of highly invasive p190bcr-abl leukemic cells. Eur. J. Cell Biol. 91: 978–987

17. De Groot RP, Raaijmakers JAM, Lammers JWJ, Jove R & Koenderman L (1999) STAT5 activation by BCR-Abl contributes to transformation of K562 leukemia cells. Blood 94: 1108–1112

18. de la Fuente MA, Sasahara Y, Calamito M, Antón IM, Elkhal A, Gallego MD, Suresh K, Siminovitch K, Ochs HD, Anderson KC, Rosen FS, Geha RS & Ramesh N (2007) WIP is a chaperone for Wiskott-Aldrich syndrome protein (WASP). Proc. Natl. Acad. Sci. U. S. A. 104: 926–931

19. Dekker FJ, Rocks O, Vartak N, Menninger S, Hedberg C, Balamurugan R, Wetzel S, Renner S, Gerauer M, Schölermann B, Rusch M, Kramer JW, Rauh D, Coates GW, Brunsveld L, Bastiaens PIH & Waldmann H (2010) Small-molecule inhibition of APT1 affects Ras localization and signaling. Nat. Chem. Biol. 6: 449–456

20. Fox M, Crafter C & Owen D (2019) The non-receptor tyrosine kinase ACK: regulatory mechanisms, signalling pathways and opportunities for attACKing cancer. Biochem. Soc. Trans. 47: 1715–1731

21. Fujimoto Y, Ochi H, Maekawa T, Abe H, Hayes CN, Kumada H, Nakamura Y & Chayama K (2011) A single nucleotide polymorphism in activated cdc42 associated tyrosine kinase 1 influences the interferon therapy in hepatitis C patients. J. Hepatol. 54: 629–639

22. Gaudet S, Branton D & Lue RA (2000) Characterization of PDZ-binding kinase, a mitotic kinase. Proc. Natl. Acad. Sci. U. S. A. 97: 5167–5172

23. Greenman C, Stephens PJ, Smith R, Dalgliesh GL, Hunter C, Bignell GR, Davies H, Teague J, Butler AP, Edkins S, Meara SO, Vastrik I, Schmidt EE, Avis T, Bhamra G, Buck G, Choudhury B, Clements J, Cole J, Dicks E, et al (2007) Patterns of somatic mutation in human cancer genomes. Nature 446: 153–158

24. Hantschel O, Warsch W, Eckelhart E, Kaupe I, Grebien F, Wagner KU, Superti-Furga G & Sexl V (2012) BCR-ABL uncouples canonical JAK2-STAT5 signaling in chronic myeloid leukemia. Nat. Chem. Biol. 8: 285–293

25. Ilaria RL & Etten RA Van (1996) P210 and P190 BCR / ABL Induce the Tyrosine Phosphorylation and DNA Binding Activity of Multiple Specific STAT Family Members. J. Biol. Chem. 271: 31704–31710

26. Jenkins C, Luty SB, Maxson JE, Eide CA, Abel ML, Togiai C, Nemecek ER, Bottomly D, McWeeney SK, Wilmot B, Loriaux M, Chang BH & Tyner JW (2018) Synthetic lethality of TNK2 inhibition in PTPN11-mutant leukemia. Sci. Signal. 11(539) eaao5617.

27. John S, Vinkemeier U, Soldaini E, Darnell JE & Leonard WJ (1999) The Significance of Tetramerization in Promoter Recruitment by Stat5. Mol. Cell. Biol. 19: 1910–1918

28. Jolma A, Yan J, Whittington T, Ukkonen E, Kivioja T & Taipale J (2013) DNA-binding Specificities of human Transcription Factors. Cell. 152(1-2) 327–339.

29. Klejman A, Schreiner SJ, Nieborowska-Skorska M, Slupianek A, Wilson M, Smithgall TE & Skorski T (2002) The Src family kinase Hck couples BCR/ABL to STAT5 activation in myeloid leukemia cells. EMBO J. 21: 5766–5774

30. Li Y, Clough N, Sun X, Yu W, Abbott BL, Hogan CJ & Dai Z (2007) Bcr-Abl induces abnormal cytoskeleton remodeling, beta1 integrin clustering and increased cell adhesion to fibronectin through the Abl interactor 1 pathway. J. Cell Sci. 120: 1436–1446

31. Li H, Yu X, Liu X, Hu P, Shen L, Zhou Y, Zhu Y, Li Z, Hui H, Guo Q & Xu J (2018) Wogonoside induces depalmitoylation and translocation of PLSCR1 and N-RAS in primary acute myeloid leukaemia cells. J. Cell. Mol. Med. 22: 2117–2130

32. Lin C, Smith ER, Takahashi H, Lai KC, Martin-Brown S, Florens L, Washburn MP, Conaway JW, Conaway RC & Shilatifard A (2010) AFF4, a component of the ELL/P-TEFb elongation complex and a shared subunit of MLL chimeras, can link transcription elongation to leukemia. Mol. Cell 37: 429–437

33. Lin, Jian-Xin, Li, Peng, Liu, Delong, Jin, Hyun Tak, He J (2012) Critical Role of STAT5 Transcription Factor Tetramerization for Cytokine Responses and Normal Immune Function. Immunity 36: 586–599

34. Lin JX, Du N, Li P, Kazemian M, Gebregiorgis T, Spolski R & Leonard WJ (2017) Critical functions for STAT5 tetramers in the maturation and survival of natural killer cells. Nat. Commun. 8(1), 1320.

35. Lionberger JM, Wilson MB & Smithgall TE (2000) Transformation of myeloid leukemia cells to cytokine independence by Bcr-Abl is suppressed by kinase-defective Hck. J. Biol. Chem. 275: 18581–18585

36. Liu J, Srinivasan S, Li C-Y, Ho I-L, Rose J, Shaheen M, Wang G, Yao W, Deem A, Bristow C, Hart T & Draetta G (2019) Pooled library screening with multiplexed Cpf1 library. Nat. Commun. 10: 3144

37. Looi CY, Sasahara Y, Watanabe Y, Satoh M, Hakozaki I, Uchiyama M, Wong WF, Du W, Uchiyama T, Kumaki S, Tsuchiya S & Kure S (2014) The open conformation of WASP regulates its nuclear localization and gene transcription in myeloid cells. Int. Immunol. 26: 341

38. Mahajan K, Lawrence HR, Lawrence NJ & Mahajan NP (2014) ACK1 tyrosine kinase interacts with histone demethylase KDM3A to regulate the mammary tumor oncogene HOXA1. J. Biol. Chem. 289: 28179–28191

39. Mahajan NP, Liu Y, Majumder S, Warren MR, Parker CE, Mohler JL, Earp HS & Whang YE (2007) Activated Cdc42-associated kinase Ack1 promotes prostate cancer progression via androgen receptor tyrosine phosphorylation. Proc. Natl. Acad. Sci. U. S. A. 104: 8438–8443

40. Mahajan K, Malla P, Lawrence HR, Chen Z, Kumar-Sinha C, Malik R, Shukla S, Kim J, Coppola D, Lawrence NJ, Mahajan NP. ACK1/TNK2 Regulates Histone H4 Tyr88-phosphorylation and AR Gene Expression in Castration-Resistant Prostate Cancer. Cancer Cell. 2017 Jun 12;31(6):790–803.e8.

41. Mahendrarajah N, Borisova ME, Reichardt S, Godmann M, Sellmer A, Mahboobi S, Haitel A, Schmid K, Kenner L, Heinzel T, Beli P & Krämer OH (2017) HSP90 is necessary for the ACK1-dependent phosphorylation of STAT1 and STAT3. Cell. Signal. 39: 9–17

42. Mahendrarajah N, Paulus R & Krämer OH (2016) Histone deacetylase inhibitors induce proteolysis of activated CDC42-associated kinase-1 in leukemic cells. J. Cancer Res. Clin. Oncol. 142: 2263–2273

43. Manser, E., Leung, T., Salihuddin, H., Tan, L., & Lim, L. (1993). A non-receptor tyrosine kinase that inhibits the GTPase activity of p21cdc42. Nature, 363(6427), 364–367.

44. Maxson, J. E., Gotlib, J., Pollyea, D. A., Fleischman, A. G., Agarwal, A., Eide, C. A., Bottomly, D., Wilmot, B., McWeeney, S. K., Tognon, C. E., Pond, J. B., Collins, R. H., Goueli, B., Oh, S. T., Deininger, M. W., Chang, B. H., Loriaux, M. M., Druker, B. J., & Tyner, J. W. (2013). Oncogenic CSF3R mutations in chronic neutrophilic leukemia and atypical CML. The New England journal of medicine, 368(19), 1781–1790.

45. Maxson, Julia., Abel, Melissa., Wang, Jinhua., Deng X., Reckel. et al (2015) Identification and Characterization of Tyrosine Kinase Nonreceptor 2 Mutations in Leukemia through Integration of Kinase Inhibitor Screening and Genomic Analysis. Cancer Res. 76: 127–138

46. McLean CY, Bristor D, Hiller M, Clarke SL, Schaar BT, Lowe CB, Wenger AM & Bejerano G (2010) GREAT improves functional interpretation of cis-regulatory regions. Nature Biotechnology. 28: 495–501

47. Menotti M, Ambrogio C, Cheong T-C, Pighi C, Mota I, Cassel SH, Compagno M, Wang Q, Dall’Olio R, Minero VG, Poggio T, Sharma GG, Patrucco E, Mastini C, Choudhari R, Pich A, Zamo A, Piva R, Giliani S, Mologni L, et al (2019) Wiskott-Aldrich syndrome protein (WASP) is a tumor suppressor in T cell lymphoma. Nat. Med. 25: 130–140

48. Meyer WKH, Riechenbach P, Schindler U, Soldaini E & Nabholz M (1997) Interaction of STAT5 dimers on two low affinity binding sites mediates interleukin 2 (IL-2) stimulation of IL-2 receptor α gene transcription. J. Biol. Chem. 272: 31821–31828

49. Mohammed, A., Zhang, C., Zhang, S., Shen, Q., Li, J., Tang, Z., & Liu, H. (2019). Inhibition of cell proliferation and migration in non-small cell lung cancer cells through the suppression of LYPLA1. Oncology Reports, 41, 973–980

50. Moriggl R, Sexl V, Kenner L, Duntsch C, Stangl K, Gingras S, Hoffmeyer A, Bauer A, Piekorz R, Wang D, Bunting KD, Wagner EF, Sonneck K, Valent P, Ihle JN & Beug H (2005) Stat5 tetramer formation is associated with leukemogenesis. Cancer Cell 7: 87–99

51. Nakken S, Gundersen S, Bernal FLM, Hovig E & Wesche J (2021) OncoEnrichR: cancer-dedicated gene set interpretation. arXiv Prepr. arXiv2107.13247

52. Negrini, S., Gorgoulis, V. G., & Halazonetis, T. D. (2010). Genomic instability--an evolving hallmark of cancer. Nature reviews. Molecular cell biology, 11(3), 220–228.

53. Nieborowska-Skorska M, Wasik MA, Slupianek A, Salomoni P, Kitamura T, Calabretta B, Skorski T. Signal transducer and activator of transcription (STAT)5 activation by BCR/ABL is dependent on intact Src homology (SH)3 and SH2 domains of BCR/ABL and is required for leukemogenesis. J Exp Med. 1999 Apr 19;189(8):1229–42.

54. Nonami A, Sattler M, Weisberg E, Liu Q, Zhang J, Patricelli MP, Christie AL, Saur AM, Kohl NE, Kung AL, Yoon H, Sim T, Gray NS & Griffin JD (2015) Identification of novel therapeutic targets in acute leukemias with NRAS mutations using a pharmacologic approach. Blood 125: 3133–3143

55. Potenta, S., Zeisberg, E., & Kalluri, R. (2008). The role of endothelial-to-mesenchymal transition in cancer progression. British journal of cancer, 99(9), 1375–1379.

56. Prieto-Echagüe V, Gucwa A, Brown DA & Miller WT (2010) Regulation of Ack1 localization and activity by the amino-terminal SAM domain. BMC Biochem. 11: 42

57. Rogakou EP, Boon C, Redon C & Bonner WM (1999) Megabase Chromatin Domains Involved in DNA Double-Strand Breaks in Vivo. J. Cell Biol. 146: 905–916

58. Sathyanarayana BK, Li P, Lin JX, Leonard WJ & Lee B (2016) Molecular models of STAT5A tetramers complexed to DNA predict relative genome-wide frequencies of the spacing between the two dimer binding motifs of the tetramer binding sites. PLoS One 11: 1–21

59. Satoh T, Kato J, Nishida K & Kaziro Y (1996) Tyrosine phosphorylation of ack in response to temperature shiftdown, hyperosmotic shock, and epidermal growth factor stimulation. FEBS Lett. 386: 230–234

60. Schafranek L, Nievergall E, Powell JA, Hiwase DK, Leclercq T, Hughes TP & White DL (2015) Sustained inhibition of STAT5, but not JAK2, is essential for TKI-induced cell death in chronic myeloid leukemia. Leukemia 29: 76–85

61. Schindler C and Darnell Jr (1995) Transcriptional responses to polypeptide ligands: The JAK-STAT pathway. Annu. Rev. Biochem. 64: 621–51

62. Shuai K, Schindler C, Prezioso V & Darnell J (1992) Activation of transcription by IFN-gamma: tyrosine phosphorylation of a 91-kD DNA binding protein. Science (80). 258: 1808–1812

63. Skene, P. J., & Henikoff, S. (2017). An efficient targeted nuclease strategy for high-resolution mapping of DNA binding sites. eLife, 6, e21856.

64. Starkova J, Madzo J, Cario G, Kalina T, Ford A, Zaliova M, Hrusak O & Trka J (2007) The Identification of (ETV6)/RUNX1-Regulated Genes in Lymphopoiesis Using Histone Deacetylase Inhibitors in ETV6/RUNX1-Positive Lymphoid Leukemic Cells. Clin. Cancer Res. 13: 1726–1735

65. Sternsdorf T, Jensen K, Reich B & Will H (1999) The Nuclear Dot Protein Sp100, Characterization of Domains Necessary for Dimerization, Subcellular Localization, and Modification by Small Ubiquitin-like Modifiers. J. Biol. Chem. 274: 12555–12566

66. Subramanian A, Tamayo P, Mootha VK, Mukherjee S, Ebert BL, Gillette MA, Paulovich A, Pomeroy SL, Golub TR, Lander ES & Mesirov JP (2005) Gene set enrichment analysis: a knowledge-based approach for interpreting genome-wide expression profiles. PNAS. 102: 15545–15550

67. Tahir R, Madugundu AK, Udainiya S, Cutler JA, Renuse S, Wang L, Pearson NA, Mitchell CJ, Mahajan N, Pandey A & Wu X (2021) Proximity-Dependent Biotinylation to Elucidate the Interactome of TNK2 Nonreceptor Tyrosine Kinase. J. Proteome Res. 20: 4566–4577

68. The ENCODE Project Consortium, Moore JE, Purcaro MJ, Pratt HE, Epstein CB, Shoresh N, Adrian J, Kawli T, Davis CA, Dobin A, Kaul R, Halow J, Van Nostrand EL, Freese P, Gorkin DU, Shen Y, He Y, Mackiewicz M, Pauli-Behn F, Williams BA, Mortazavi A, et al (2020) Expanded encyclopaedias of DNA elements in the human and mouse genomes. Nature 583: 699–710

69. Thomas-Chollier M, Herrmann C, Defrance M, Sand O, Thieffry D & van Helden J (2011) RSAT peak-motifs: motif analysis in full-size ChIP-seq datasets. Nucleic Acids Res. 40: e31–e31

70. Tormo AJ & Gauchat J-F (2013) A novel role for STAT5 in DC: Controlling the Th2-response. JAK-STAT 2: e25352–e25352

71. Toscano MG, Anderson P, Muñoz P, Lucena G, Cobo M, Benabdellah K, Gregory PD, Holmes MC & Martin F (2013) Use of zinc-finger nucleases to knock out the WAS gene in K562 cells: a human cellular model for Wiskott-Aldrich syndrome. Dis. Model. Mech. 6: 544–554

72. Uchida E, Suwa S, Yoshimoto R, Watanabe K, Kasama T, Miura O & Fukuda T (2019) TOPK is regulated by PP2A and BCR/ABL in leukemia and enhances cell proliferation. Int J Oncol 54: 1785–1796

73. van der Horst EH, Degenhardt YY, Strelow A, Slavin A, Chinn L, Orf J, Rong M, Li S, See L-H, Nguyen KQC, Hoey T, Wesche H & Powers S (2005) Metastatic properties and genomic amplification of the tyrosine kinase gene ACK1. Proc. Natl. Acad. Sci. 102: 15901–15906

74. Wang A, Pei J, Shuai W, Lin C, Feng L, Wang Y, Lin F, Ouyang L & Wang G (2021) Small Molecules Targeting Activated Cdc42-Associated Kinase 1 (ACK1/TNK2) for the Treatment of Cancers. J. Med. Chem. 64: 16328–16348

75. Wang C, Sample KM, Gajendran B, Kapranov P, Liu W, Hu A, Zacksenhaus E, Li Y, Hao X & Ben-David Y (2021) FLI1 Induces Megakaryopoiesis Gene Expression Through WAS/WIP-Dependent and Independent Mechanisms; Implications for Wiskott-Aldrich Syndrome. Front. Immunol. 12:607836

76. Warmuth M, Bergmann M, Prieß A, Häuslmann K, Emmerich B & Hallek M (1997) The Src family kinase Hck interacts with Bcr-Abl by a kinase-independent mechanism and phosphorylates the Grb2-binding site of Bcr. J. Biol. Chem. 272: 33260–33270

77. Weber A, Borghouts C, Brendel C, Moriggl R, Delis N, Brill B, Vafaizadeh V & Groner B (2015) Stat5 exerts distinct, vital functions in the cytoplasm and nucleus of Bcr-Abl+ K562 and Jak2(V617F)+ HEL leukemia cells. Cancers (Basel). 7: 503–537

78. Xie S, Wang Y, Liu J, Sun T, Wilson MB, Smithgall TE & Arlinghaus RB (2001) Involvement of Jak2 tyrosine phosphorylation in Bcr-Abl transformation. Oncogene 20: 6188–6195

79. Yokoyama N & Miller WT (2003) Biochemical properties of the Cdc42-associated tyrosine kinase ACK1. Substrate specificity, autophosphorylation, and interaction with Hck. J. Biol. Chem. 278: 47713–47723

80. Yokoyama N, Lougheed J & Miller WT (2005) Phosphorylation of WASP by the Cdc42-associated kinase ACK1: Dual hydroxyamino acid specificity in a tyrosine kinase. J. Biol. Chem. 280: 42219–42226

81. Xu, X., Sun, Y. L., & Hoey, T. (1996). Cooperative DNA binding and sequence-selective recognition conferred by the STAT amino-terminal domain. Science 273(5276), 794–797.

82. Zhang H, Wang Y, Yang H, Huang Z, Wang X & Feng W (2021) TCF7 knockdown inhibits the imatinib resistance of chronic myeloid leukemia K562/G01 cells by neutralizing the Wnt/β-catenin/TCF7/ABC transporter signaling axis. Oncol Rep 45: 557–568

83. Zhang Q, Wang HY, Wei F, Liu X, Paterson JC, Roy D, Mihova D, Woetmann A, Ptasznik A, Odum N, Schuster SJ, Marafioti T, Riley JL & Wasik MA (2014) Cutaneous T Cell Lymphoma Expresses Immunosuppressive CD80 (B7-1) Cell Surface Protein in a STAT5-Dependent Manner. J. Immunol. 192: 2913 LP –2919

84. Zhang Y, Lee C, Geng S & Li L (2019) Enhanced tumor immune surveillance through neutrophil reprogramming due to Tollip deficiency. JCI Insight 4 (2) e122939.

85. Zhu F, Zykova TA, Kang BS, Wang Z, Ebeling MC, Abe Y, Ma W, Bode AM & Dong Z (2007) Bidirectional Signals Transduced by TOPK-ERK Interaction Increase Tumorigenesis of HCT116 Colorectal Cancer Cells. Gastroenterology 133: 219–231

86. Zigmond SH (2000) How WASP regulates actin polymerization. J. Cell Biol. 150: F117–F120

